# Non-Local Conceptual Combination

**DOI:** 10.1101/2022.12.11.519989

**Authors:** Alicia Parrish, Amilleah Rodriguez, Liina Pylkkänen

**Author notes:** Corresponding author: Alicia Parrish.

## Abstract

It is uncontroversial that the syntax of an expression largely determines its meaning. For example, there is no way to interpret a sentence like “the blue hat has a white bow” as telling you that there is a white hat that has blue bow. But to what extent are the brain’s combinatory interpretive routines exclusively locked into the structures given by syntax? Consider another example: “The blue color of his hat is pretty.” This sentence tells us that a color is pretty, that the color is blue and that the color belongs to a hat. What the syntax of this sentence does not give us is a combination of “blue” and “hat.” But clearly, if we were to draw a picture of the meaning of this sentence, it would have a blue hat in it. We asked: upon encountering “hat” in this sentence, do our brains combine the features of “blue” with the features of “hat,” despite the long distance between them and no direct syntactic relation? By using a known neural measure of conceptual combination in the left anterior temporal lobe, we obtained evidence using MEG that our brains appear to perform such a long-distance conceptual combination that does not track the syntax. Intriguingly, word (or rather concept) order affected the directionality of the effect. While the effect of concept order remains a topic for future work, our results overall suggest that the meaning composition system of language is likely richer than the combinatory steps predicted from syntactic structures.

## INTRODUCTION

A key component of language is the ability to compose meaning from words not linearly adjacent. Long-distance dependencies have been well-studied in constructions involving displacement of an element in the syntax, as in questions or relative clauses, but in this study, we are interested in a different type of relation: elements that conceptually combine across a long-distance. Consider the phrase “a blue hat” versus the phrase “a hat that is a lovely shade of blue.” Both result in a conceptual representation that contains a blue hat, even though “blue” and “hat” locally compose only in the former case. The neurobiology of language has not yet probed such expressions. But they offer an interesting test case, both for cognitive and neurobiological theories of languages: evidence for a conceptual tier of combinatory processing that does not track syntax should inform our theories of the syntax-semantics interface, and uncovering whether a neural site of conceptual combination is able to operate across a long-distance would critically inform the mechanism underlying the measured activity. In the current work, we tested whether a well-studied neural correlate of conceptual combination, localized in the left anterior temporal lobe (LATL), shows sensitivity to long-distance conceptual combination. To further probe the underlying mechanism, we also performed a decoding analysis to track the activation of the first element across the intervening words before the second element was encountered, the logic being that such maintenance should occur in order for the two concepts to compose later.

We build on a large body of work showing that LATL activity reflects conceptual as opposed to syntactic aspects of composition, both in minimal phrases (e.g., Baron and Osherson, 2011; Baron et al., 2010; Blanco-Elorrieta and Pylkkänen, 2016; Kim and Pylkkänen, 2019; Westerlund and Pylkkänen, 2014; Zhang and Pylkkänen, 2015, 2018a,b) and in full sentences (Poortman and Pylkkänen, 2016; Parrish & Pylkkänen, 2022; Kim & Pylkkänen, 2021; Flick & Pylkkänen, 2020; Zhang & Pylkkänen, 2018). Though these studies have substantially advanced our understanding of the LATL’s role in language processing, we do not know if the LATL can operate across a long distance when there is no direct syntactic relation between the two composing concepts. In other words, does the LATL reflect a truly general combinatory operation that is *independent* of syntactic structure? Or a local combinatory operation, closely tracking syntax? We address this question using magneto-encephalography (MEG) by comparing structurally-matched sentences that vary in whether two words in separate syntactic phrases combine to form a single conceptual representation (as “blue” and “hat” do in “the blue color of this hat is pretty”) or do not combine into a single conceptual representation (“blue” and “hat” do not conceptually compose in “the blue lamp near this hat is pretty”).

We will use the term “LATL conceptual combination” to refer to conceptual combination as it is reflected by LATL activity peaking at 200-300ms after the onset of a word in a combinatory context. We do not mean to imply that aspects of conceptual combination might not be implemented elsewhere in the brain as well, rather, it is likely that this is the case (Coutanche et al., 2019; Price et al., 2015, i.a.). For the LATL, the original finding from Bemis and Pylkkänen (2011) showed that, measuring on the target word “boat,” there was greater activation in the LATL when “boat” was preceded by “red” (for the combinatory context “red boat”) as opposed to (i) a consonant string like “xkc,” and (ii) when it was presented in a list context, following another noun, like “cup.” The LATL has been hypothesized to be a crucial component of humans’ combinatory capacity in language due not only to these studies in healthy adults, but also based on language deficit studies (e.g., Lukic et al., 2021; Mesulam et al., 2019, 2015; Wilson et al., 2014). It is broadly accepted that the LATL is an important region for language tasks, and it has been posited to be a general “semantic hub” (Lambon Ralph et al., 2017).

A substantial body of prior literature on the LATL’s role in composition has already shed light on the mechanism underlying LATL’s composition sensitivity. In addition to the original findings that two-word phrases elicit more LATL activity than two-word lists in both comprehension (Bemis and Pylkkänen, 2011) and production (Pylkkänen et al., 2014), MEG studies have also shown an increase in LATL activity not only for adjective-noun phrases, but also within a noun-noun phrase (Flick et al., 2018), adverb-verb phrase (Kim and Pylkkänen, 2019), verb-argument phrase (Westerlund et al., 2015), and adverb-adjective phrase (Parrish and Pylkkänen, 2022). Despite this generality, different conceptual or featural aspects of the words being combined affects the LATL’s response. For example, Zhang and Pylkkänen (2015) compared phrases where they contrasted whether the modifier or the head of a phrase was more/less specific; for example “tomato dish” has a more specific modifier than “vegetable dish,” and “tomato soup” has a more specific head noun than “tomato dish.” They found that making an already specific concept (“soup”) even more specific led to lower LATL activation compared to modifying a less specific concept (“dish”). This study and related ones showed that the observed increase in LATL activation is not due to syntactic composition, but rather indexes, at least to some extent, incremental changes in the featural specificity of words (Westerlund and Pylkkänen, 2014; Zhang & Pylkkänen, 2018; Kim & Pylkkänen, 2021). In sum, we already know that the LATL’s response to words in combinatory contexts is a general process in terms of modality and word category, and we know that composition in the LATL is sensitive to conceptual information rather than syntactic information. Prior work has even shown that LATL conceptual combination can operate on two adjacent concepts even if they do not locally merge in the syntax (Parrish & Pylkkänen, 2022). The current study tests if such syntax-independent composition can occur across a long distance.

As regards non-grammatical factors modulating the LATL, it has also shown sensitivity to level of association between the composing words, with high-association pairs (“French cheese”) driving LATL activity more than low association pairs (“Korean cheese”) (Li and Pylkkänen, 2021). Parrish and Pylkkänen (2022) also showed that plausible modifications lead to a greater increase in LATL activity relative to implausible modifications. This finding was surprising since plausibility cannot be known until a meaning has been constructed. Thus the LATL conceptual combination effect should be too early in the processing stream to reflect plausibility. However, plausible combinations are also more highly associated than implausible ones. Thus we conjecture that the plausibility finding of Parrish and Pylkkänen (2022) and the association finding of Li and Pylkkänen (2021) may both be instances of increased combinatory activity for high association composition.

Pertaining to the relation between composition and the grammar, Mollica et al. (2020) found that only fairly local combinations engage the language network when stimuli are ungrammatical. They defined local as pertaining to two words that are close enough to enter into a dependency relation, and they showed that local perturbations in word order did not affect the BOLD signal measured, relative to fully grammatical sentences. Taken together, these studies raise the question of how incremental (i.e., word-by-word) changes in the brain’s feature space representations interact with composition, and these studies highlight the need to more fully separate or control for these effects when studying the parser’s underlying combinatory capacity. With this in mind, we attempt to block this automatic sensitivity to local changes by separating the composing items with intervening lexical material, specifically making sure there is at least one intervening content word.

### Two hypotheses about LATL conceptual combination: generalized vs. locally-dependent composition

Our study aims to distinguish between two competing hypotheses about the mechanism of LATL conceptual combination. Under the first mechanism, which we will call *generalized composition*, the LATL reflects all aspects of conceptual combination that contribute to sentence meaning, whether those instances of conceptual combination map onto instances of syntactic composition or not. If two concepts are combined in the composite meaning of the sentence, the LATL created that combination. In opposition, the second hypothesis proposes that what has been referred to as LATL conceptual composition in the literature of the past about ten years reflects a local operation. We call this *locally-dependent composition*. As the critical test, we separate the two words that are in combinatory or non-combinatory contexts with intervening lexical material and place the color concept in a syntactic position from which it cannot directly predicate of the noun.

The predictions of the opposing hypotheses are diagrammed in Figure 1.C.i. Generalized composition predicts long-distance conceptual combination to show the same contrast between Combinatory and Non-combinatory contexts as a Local control conditions, in which the composing words are linearly adjacent. In contrast, locally-dependent composition predicts an effect of LATL composition only for the Local controls.

**Figure 1:**
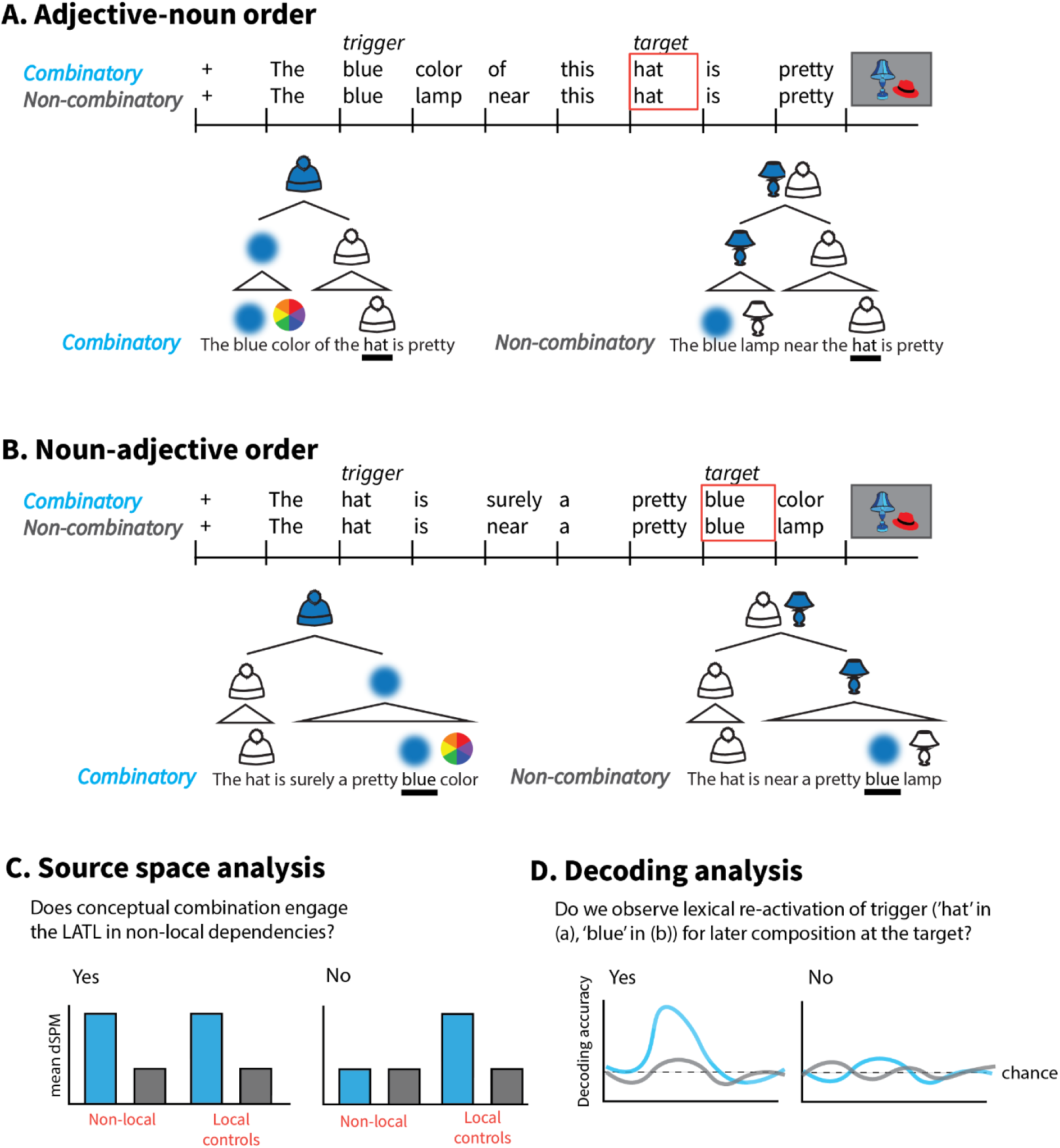
Possible patterns of results from the planned analyses for both the Adjective-Noun order stimuli (A) and the Noun-Adjective order stimuli (B). Within the two word order conditions, the Combinatory and Non-combinatory structures are broadly identical. (C) shows the predictions relevant for analyses conducted on the target word in the spatiotemporal source space analysis. (D) shows the predictions for the decoding analysis.

### Decoding word representations

To preview the results, we ultimately find evidence in favor of the generalized composition hypothesis, indicating that LATL conceptual combination reflects a process that is more general than just minimal phrasal composition and, in fact, does not rely on linear adjacency or having the two combining words be in the same phrase. The question that then emerges is what representations conceptual combination is accessing for these results to hold. We focus next on the question of how concepts become available to the LATL for combination by asking whether we find evidence of the reactivation of lexical items in some sentential contexts but not others. We use a decoding analysis to investigate when specific lexical representations from early in the sentence are available for composition at different points later in a sentence. We consider whether lexical re-activation of the earlier word at the point of composition (i.e., reactivating “blue” at the final word in “the blue color of this hat”) is a potential explanation for how conceptual combination can selectively engage for some sentences and not others, even when those sentences are matched for syntactic structure.

Two of the previous studies that investigated adjective-noun composition using decoding analyses (Fyshe et al., 2019; Honari-Jahromi et al., 2021) have suggested that conceptual combination may act on two co-activated lexical representations. Fyshe et al. (2019) found that the semantic features of the adjective were decodable during presentation of the noun (see also Fyshe, 2015), but they were unable to determine if that extended decodability was due to a re-emergence of the adjective’s lexical representation for composition, as opposed to sustained or residual activation of those features. These two possibilities were not distinguishable in their study because the adjective and noun were linearly adjacent. If co-activation of the modifier and modified concept is required for composition, then we would expect to be able to observe evidence of a lexical representation being reactivated at the point of composition, even when those words are non-adjacent.

In a naming task, Honari-Jahromi et al. (2021) found an increase in decodability of nouns in combinatory tasks (naming the object on a screen and its color, e.g., “green lamp”) compared to a list task (naming the object on a screen and the background color of the image, e.g., “green, lamp”), opening up the possibility that the “activeness” of lexical representations is tied to the ways that the brain will need to use those representations, rather than just whether they need to become activated at all. This study also identified an asymmetry in the decodability of adjective vs. noun representations, such that the nouns were more robustly decodable over a longer time period compared to adjectives, even when both were part of a combinatory context. In our study, we compare Combinatory and non-Combinatory contexts in sentences where the representations being decoded occur earlier in the sentence and need to be remembered (but not yet composed) in order to complete the task, a design that will allow us to further probe how these representations are sensitive to the way they will be later engaged. In order to make sure we are measuring reactivated concepts rather than residual activation from a previous word, we separate the composing (or non-composing) words by intervening lexical material. The current study is well-positioned to shed light on whether co-activation of two lexical representations is a likely pre-requisite for combinatory operations, and whether the previous findings from Fyshe et al. (2019) are the result of residual activation from the modifier, or the reactivation of the modifier at the noun.

This paradigm in which we ask which earlier words are being activated at which points of a sentence will likely make readers think of other kinds of non-local dependencies, such as filler-gap kinds of constructions. Language comprehension often requires making a link to a phrase (or concept) that appeared earlier in the sentence or discourse. In English, this is the case with pronominal referents (e.g. “I saw a toy on the ground but then a girl picked it up” requires linking “it” with “a toy”), relative clause structures (“The dog that I love to kiss ran away” requires understanding “the dog” as the patient of “kiss”), cleft structures (“It was the lizard that Suzie took a picture of yesterday” requires interpreting “the lizard” in the gap site), and *wh*-dependencies (“Who did Jackie say will call you?” requires interpreting the wh-word as the agent of “call”), among others. In all these cases, meaning composition must occur between words that are not linearly adjacent, and often between words that are in different clauses.

Most debates about how humans come to understand these kinds of non-contiguous sequences have centered around how syntactic or semantic composition works for different dependencies, but the same questions apply when asking how conceptual combination works in these types of constructions. Unlike with syntactic composition, though, we have a well-defined MEG component that allows us to measure effects of conceptual combination. Rather than building contrasts that attempt to isolate syntactic effects, we can use matched syntactic constructions to create contrasts that we know are relevant for conceptual combination—which is precisely what we have done in this study. Although investigating combinatory processing in other kinds of non-local dependencies through similar measures as we propose here would be valuable, it would ultimately be harder to connect to the broader literature on syntactic composition, as there is no single effect that consistently indexes certain types of syntactic operations.

Specifically regarding conceptual combination, our study asks whether (i) meaning-based cues trigger the reactivation of a specific lexical item from earlier in the sentence, and whether (ii) the re-activated concepts are sufficiently rich to be decodable at that later point in time. Although previous studies of word reactivation in long distance dependencies, particularly those that use filler-gap constructions, have looked at activation in the LIFG as a measure of structural complexity and processing load (Stromswold et al., 1996, i.a.), increases in activation can depend on whether there is similarity-based interference from another noun, and the timecourse of activation can vary with different filler-gap constructions (Leiken et al., 2015).

We use a decoding analysis, as opposed to looking at LIFG activation patterns as many other studies have, to answer this question to investigate whether the actual lexical item gets reactivated as opposed to whether one of the sentence structures is more or less difficult. Answering these questions will lay the groundwork for determining how conceptual combination interacts with syntactic parsing operations, specifically by narrowing the space of possibilities of what kinds of representations the mechanism needs to have access to. Figure 1.C.ii lays out the two hypotheses associated with determining whether re-activated concepts are decodable at the point when they need to compose.

## METHODS

### Participants

We recruited 22 right-handed participants for this study. All participants reported having learned English at an early age, and they reported no history of language-related or general cognitive impairments and had normal (or corrected-to-normal) color vision. We set a cutoff of 80% overall accuracy on the task as inclusion criteria for each participant (chance rate was 50%; see task details and full behavioral results are described in later sections). Based on this threshold, no participants needed to be removed from analysis, and we assume that all were adequately paying attention throughout the entire task and that they understood the content of the stimuli.

### Materials

All stimuli in this study were created by the experimenters. Sentence stimuli were created through the use of templates with different words slotted in, and images for the task were created using freely-available vector images manipulated with Adobe Illustrator. This section describes the creation of all stimuli as well as the procedure used to ensure that all sentences used in the experiment were as natural as possible.

### Stimuli creation

We created a small vocabulary made up of four target nouns and four target color adjectives. The small vocabulary was necessary to ensure that, in the decoding analysis, there were enough trials with each of the target words to be able to acquire enough signal in the sensor data to train a classifier. To ensure some lexical diversity and make sure the sentences were not overly repetitive with only the target words varying, the vocabulary also consisted of five different position words, three verbs of seeing, five adverbs, five adjectives, four intensifiers, five color descriptors, four words for screens, four non-target colors, and seven non-target nouns. The full vocabulary is provided in Appendix Table 5, and words from that vocabulary were slotted in to the templates specific to each condition, as shown in Appendix Table 6. The target nouns and target color words were systematically slotted in to create all 16 pairing possibilities in equal proportion, and we ensured that no sentence contained two of the same vocabulary words. The additional vocabulary items that were included for lexical diversity were selected randomly for each template but kept fully consistent within a single noun+color pairing to ensure that the critical experimental comparisons were fully matched for the lexical properties of the additional vocabulary.

The target nouns were selected to be easily imageable and to be maximally distinct from the other potential target nouns that were considered, as measured by cosine similarity of the words’ word2vec (Mikolov et al., 2013) embeddings. The target colors were selected to be common primary or secondary colors (plus black) that are distinct from each other on a computer screen and reasonable colors for all the target nouns to be (i.e., they’re not independently implausible as modifiers of any of the target nouns).

### Stimuli norming

Stimuli selection. Because stimulus sentences were created through a template, we verified that the stimuli were natural-sounding and semantically plausible to avoid creating any additional processing load. For example, “the key is above the chair” is much more “natural” than “the chair is above the key” (as verified in our stimulus norming). We selected all possible combinations of two target nouns in different positions with each other (e.g., for the nouns “lamp” and “hat,” we got judgments for sentences that included “the lamp is above the hat,” “the hat is above the lamp,” “the lamp is near the hat,” etc.). This led to 100 unique combinations (5 target nouns * 4 different target nouns * 5 position words). We also included a total of 125 sentences from the other 6 experimental conditions that did not include a position word to ensure that these sentences were also reasonably acceptable.

Participants judged the sentences along a 7-point scale, with 1 defined as “very unnatural” and 7 defined as “very natural.” We instruct participants to answer in accordance to what sounds typical or grammatical to them, as we are looking for subjective judgments about the plausibility and interpretability of these items. The exact wording of the instructions and the examples provided to participants are listed in Appendix B. Each participant rated 21 sentences, of which 15 were experimental items and 6 were attention checks to ensure that participants were paying attention and using the whole 7-point slider.

We recruited participants on Amazon Mechanical Turk (MTurk). When using a crowdsourcing platform for an experimental study, it is important to ensure that the participants are felicitously attempting the task. For this reason, we employed fillers that served as attention check items, using both instructed-response items and gold-labeled items (Pei et al., 2020; Sheehan, 2018). For half of the fillers, the example was an instructed-response item, meaning that the example explicitly told the participant what number they needed to select on the scale (e.g., “The option to select is seven on this item.”), with one number towards the very natural side, one number towards the very unnatural side, and one number in the middle. For the other half of the filler items, we used gold-labeled examples. The gold-labeled examples were sentences that we wrote to be intentionally natural (e.g., “Those purple toys are laying on the floor”), intentionally unnatural for semantic reasons (e.g., “Those dogs have black spots but those same dogs do not”), or intentionally unnatural due to the position word (“The floor is laying on those purple toys”), and so we had already annotated the expected response. All the attention check items are provided in Appendix B, along with total accuracy rates on those items from the participants.

On MTurk, we made the task open to all workers located in the US with at least 5000 tasks (HITs) already completed and with a task approval rating at or above 99%, both standard filters used in MTurk studies to ensure better quality of results (Daniel et al., 2018). A total of 279 participants completed the task. The task took approximately 3 minutes to complete, and we paid participants 75 cents, so long as they passed at least two out of three instructed response fillers (attention check items that name an explicit number to answer; 7 participants got fewer than 2 of these items correct, and so we rejected their work, as they were likely answering at random without reading the examples). For their data to be included in our analysis, participants had to answer within the expected ranges on at least five out of six (83.3% accuracy) of the attention check items. A total of 226 participants (83.1%) passed this additional threshold, and their data was included in our analysis.

Each sentence was rated by at least 18 unique participants. We consider average scores above 4 on this scale to indicate that a sentence was acceptable. We computed how often a noun in each position (either as the first noun or the second noun) was rated as being acceptable. The results of this analysis are in Table 4 of Appendix B. We excluded from the final stimuli any cases where an object was not rated as natural in that position at least half the time (e.g., “book” was always rated as natural when it was *above* something, but only rated as natural 25% of the time when it was *below* something, so we excluded all sentences where a book is below another object). We did this so that objects only ever appeared in reasonably canonical positions with regard to other objects, and to reduce the chances that a participant will be able to predict the upcoming noun, given our fairly small set of vocabulary. We made sure that there were multiple reasonable continuations of the sentence after the presentation of the position word. To also make sure that all of the final stimuli were natural, we additionally excluded any pairings that were idiosyncratically rated low, such as sentences like “the chair is under the book”, where on their own, chairs being under an object was acceptable more than half the time, and something being under a book was acceptable more than half the time, but the combination of those elements in this sentence was rated as very unnatural.

The crucial comparison for the norming study was between Combinatory and Non-Combinatory conditions within each word order and locality condition. These results are summarized in Table 1 and plotted in Appendix Figure 8. There was no significant difference of Combination in the Non-local Noun-Adjective order stimuli (t = 0.30, p = 0.77); however, this comparison was significantly different for the Adjective-Noun order stimuli (t = 6.49, p < 0.01), likely due to the very low score for sentences in the Local Non-Combinatory condition.

**Table 1:** Results from the stimuli norming study. ‘Mean score’ represents the mean of the likert-scale judgments made by participants. “Adj.” is short for “Adjective”

| Word order | Locality | Combination | Mean Score |
| --- | --- | --- | --- |
| Adj. -<br>Noun | Non - Local | Combinatory | 6.1 |
|  |  | Non-Comb. | 4.4 |
|  | Local | Combinatory | 5.4 |
|  |  | Non-Comb. | 2.9 |
| Noun -<br>Adj. | Non - Local | Combinatory | 5.3 |
|  |  | Non-Comb. | 4.7 |
|  | Local | Combinatory | 6.1 |
|  |  | Non-Comb. | 5.7 |

Given the need for the stimuli to be balanced in terms of word position and combinatory context, this difference in average acceptability of the stimuli was unavoidable. As the average acceptability of the crucial Non-local condition was still above the scale’s midpoint of 4 on a 7-point scale (and mostly above 4.5), we point this out as a potential source of difference, but one that is not likely to have a strong impact on the results. In fact, the lower acceptability of the Non-combinatory stimuli in this case would, if anything, result in greater activation compared to the more acceptable Combinatory stimuli, pushing the effects in the opposite direction of what we hypothesize if LATL conceptual combination effects obtain in non-local dependencies.

Within the Local controls, there was a more striking difference between conditions, with the Non-combinatory stimuli in the Adjective-Noun order condition being rated much lower than all other conditions. This difference is likely due to the very high difficulty of creating a sentential context where an adjective and noun are adjacent (or nearly adjacent), and yet they do not conceptually compose. This condition relied on gapping to achieve this difference (e.g., “This screen’s lamps are blue, hats red, and socks gray”). Due to the need for highly controlled stimuli, we considered this difference unavoidable, but potentially important when interpreting the results, and we will return to this in the discussion section.

### Experiment design

This experiment used a 2 x 2 x 2 design of Word order (Noun-Adjective, Adjective-Noun) by Locality (Local, Non-local) by Combination (Combinatory, Non-combinatory). The relevant contrasts for each of these conditions is laid out in Table 2. The two Word order conditions, Adjective-Noun and Noun-Adjective, refer to the order of the two critical (bolded) words, and reflects the order in which they appear in the sentence. The two Locality condition, Non-local and Local (controls), refer to whether the two critical words were separated by intervening content words. In the Non-local condition, there were always three or four words between the two critical words (the trigger word and the target word), and at least one of those intervening words was a content word (i.e., it is not a determiner, preposition, or auxiliary verb). In the Local condition, the two critical words are either adjacent or only separated by the word is or a. Finally, Combination (Combinatory, Non-combinatory) refers to whether the context for the two critical words is one in which they will conceptually compose or not. In the Combinatory condition, the two critical words will compose to form a more specific concept—a blue hat in the examples in Table 2. In the Non-combinatory context, it is not necessarily the case that there is no composition, but rather there is no conceptual composition between the trigger word and the target word—that is, upon hearing a sentence in the Non-combinatory condition, the person would not need to build the conceptual representation of a blue hat to understand the sentence.

**Table 2:**
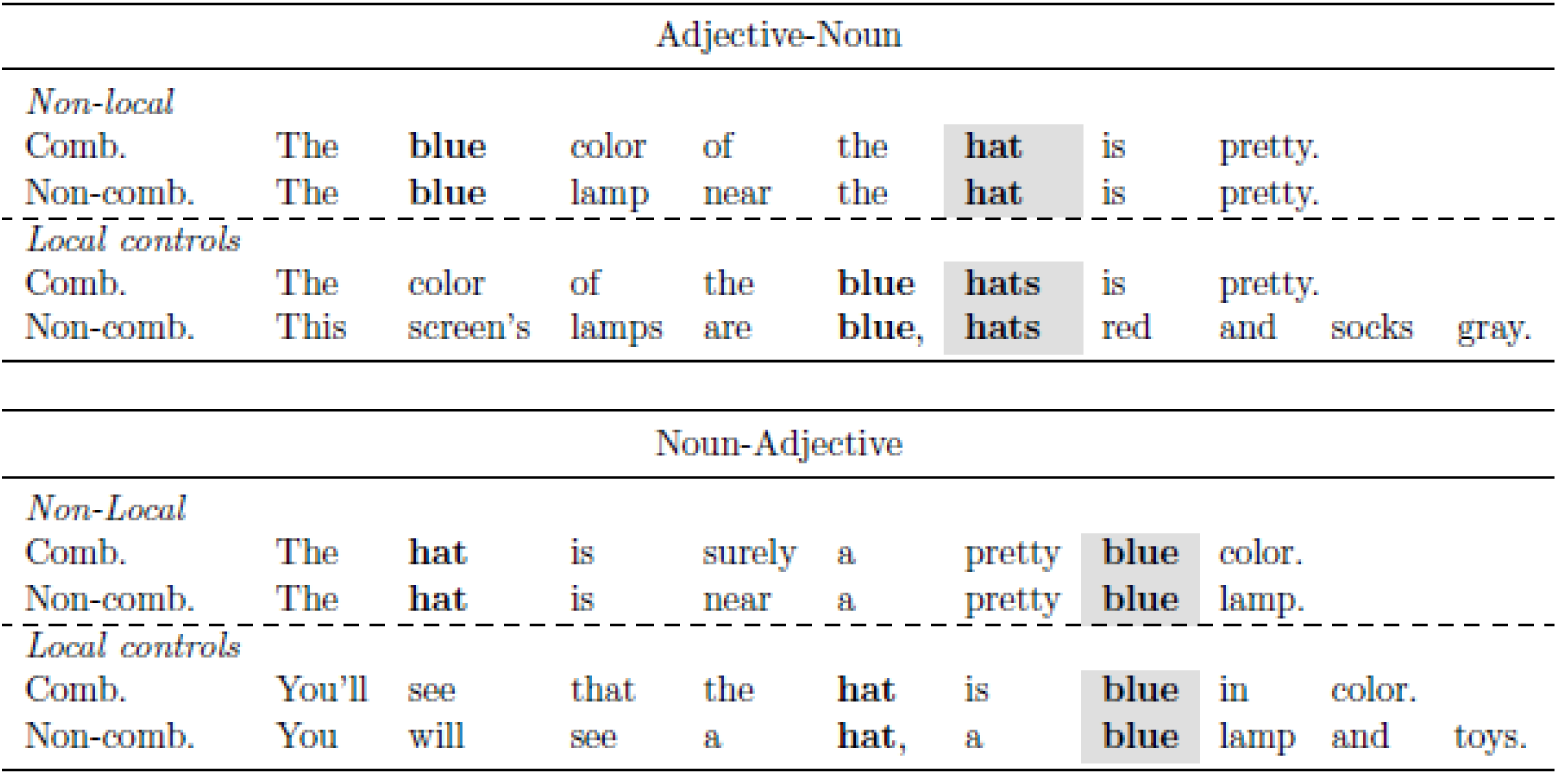
Example stimuli from Experiment 2. The target word is shaded for each condition. In Adjective-Noun order stimuli, the target word is always in the position of “hat” (the 6th word), and in Noun-Adjective order stimuli, the target word is always in the position of “blue” (the 7th word). The first bolded words in each example sentence represents the trigger word. The main contrast in whether that trigger word conceptually composes at the point of the shaded target word. In the Combinatory examples, they do conceptually compose, and in the Non-combinatory examples, they do not.

As another important point, there was also no direct syntactic relationship between the two words that conceptually composed in the Non-local examples. This relationship is schematized in Figure 1, which shows the assumed relationship between the conceptual and syntactic structure as it is being built. Crucially, in all cases, the Non-local sentences had the trigger word and the target word in different syntactic phrases, so they were separated both linearly and structurally.

The Noun-Adjective word order examples were part of a construction that has not been well studied in terms of conceptual combination yet. Both Combinatory conditions in the Noun-Adjective word order relied on a predication structure to achieve an analogous conceptual representation to what the Adjective-Noun condition can do with minimal phrasal composition in the Local controls. As discussed in the introduction, there is some evidence that Noun-Adjective word order should not matter for the LATL for combinatory effects and that words in combinatory predication structures can also drive LATL activity relative to non-combinatory controls. The possibility of making an even more broad and generalizable claim about the way that LATL conceptual combination works makes the addition of this condition worthwhile, but we note that it is not as well vetted a structure for this kind of paradigm as the examples in the Adjective-Noun word order condition.

### Procedure

Following the guidelines set out by our Institutional Review Board, all participants gave informed, written consent before beginning the study. We employed two slightly different tasks. Participants first completed a short single-word picture-verification task (described in the appendix), and then they completed the main task, which was a picture-verification task with full sentences as stimuli. The main task was split into two halves, so that all the Adjective-Noun order stimuli were together in one half and all of the Noun-Adjective order stimuli were in the other half. Trial presentation within each of those halves was fully randomized. The order of presentation of these two halves was counterbalanced across all participants.

We measured MEG responses during sentence reading using RSVP in a picture-verification task. Stimuli were presented visually, one word at a time, via PsychoPy (Peirce, 2007). Each trial began with a fixation cross at the center of the screen that lasted for 500ms, followed by a 150ms blank screen. The words were presented with a 450ms SOA (150ms ISI). All sentences were eight to ten words in length and contained normal English capitalization of the first word and normal punctuation (commas, periods) throughout.

On one third of the trials, the sentence was followed with a picture-verification task. Which trials were followed by a task, and which were not followed by a task was determined randomly, and the presentation order of task and non-task trials was also fully randomized. The purpose of this task was to assess participants’ comprehension of the stimuli and to ensure that they paid attention throughout the study. In general sentence understanding, we assume that composition is required, and that hearing a full sentence may be sufficient to push the parser into a composition task mode. In this sense, composition is automatic provided the participant is paying attention to the sentence, so we used a task that ensures attention. Further, to make sure that composition is required for the task, we ensured that correctly responding to the task required participants to pay attention to potentially all aspects of the sentence: the name of the object(s), the color of the object(s), and/or the relative position of the object(s).

Figure 2 shows an example of a single trial when it was followed by a task. In this example, the correct answer is that the image does not match the sentence, as there is no blue hat pictured near a lamp. On trials where there was no task, the screen following the last word of the sentence still asked, “Ready for next?” to ensure that participants were ready for the next sentence to begin and to allow participants to control the pacing of the experiment for themselves. The entire task took approximately one hour and was segmented into 12 blocks to allow participants to take a short break every few minutes to stretch or to rest their eyes. All breaks were self-paced, and participants saw feedback about their overall accuracy.

**Figure 2:**
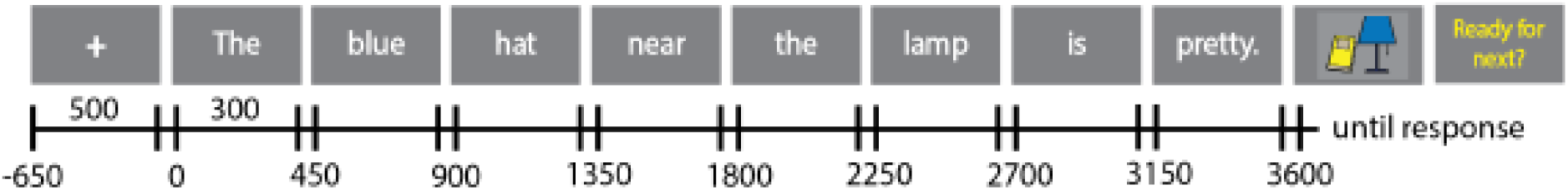
Diagram of the procedure used for the main picture-verification task. Only 33% of the sentences were followed by a picture. In cases where there was no task, participants saw the screen that asks, “Ready for next?” after the end of the sentence.

### Data acquisition

We digitized participants’ headshapes using the Polhemus FastSCAN system (Polhemus Inc., Colchester, USA). This headshape data, along with fiducial landmarks, was co-registered to the average brain available in FreeSurfer (Dale et al., 1999). Continuous MEG data was recorded using a 208-channel axial gradiometer whole-head system using a 1000Hz sampling rate. All preprocessing was done using MNE version 1.0.1 (Gramfort et al., 2014) and eelbrain version 0.37 (Brodbeck et al., 2021, 2022). The MEG data was recorded with a 200 Hz low pass filter and 0.1 Hz high pass filter, and then filtered offline using a Maxwell spatial filter followed by a 1-40 Hz bandpass filter after identifying and interpolating excessively noisy, saturated, or dead MEG channels.

### Processing of MEG source data

Initial artifact rejection was completed with independent component analysis (ICA) of the filtered data for each participant to remove artifacts due to known environmental noise, eyeblinks, and heartbeat, as is standard for MEG studies. During data processing, we found that one participant had excessive environmental noise during their recording that resulted significant data loss and over 20% of the sensors becoming saturated by the end of the experiment. We removed this participant from all analyses.

Epochs were created starting from 100ms before the start of the sentence trial and extending to 50ms after the end of the target word in the Noun-Adjective condition (from −100ms to 3200ms from sentence onset) to capture changes throughout the entire sentence and to allow for baseline correction in the window before the beginning of the sentence. Epoch rejection was done via an absolute threshold to eliminate epochs containing amplitudes exceeding 2e-12 fT and a peak-to-peak threshold to eliminate epochs containing amplitude changes greater than 3e-12 fT, as both these cases were assumed to contain artifacts not removed with ICA. In total, 0.7% of all epochs were rejected, with no individual participant having greater than 5% of their trials rejected. Two participants had to end the study early, resulting in one losing about a quarter of trials from the Noun-Adjective order condition and the other losing about a quarter of trials from the Adjective-Noun order condition. For the half of the experiment that each subject was missing data from, we removed that participant from the decoding analysis. In the source space analysis, the total number evoked responses in each condition were equalized for each participant.

We applied baseline correction to the period of each epoch starting from 100ms before the onset of the first word of the sentence trial to the start of that trial. Because the target words were in the middle of a sentence, we could not apply baseline correction directly before the target word as this time region (i) captures active language processing and (ii) is very likely to encode difference already present between the experimental conditions. Doing baseline correction at the beginning of the sentence also allowed us to expand post-hoc exploratory analyses to earlier regions in the stimuli. Channel noise covariance was estimated from the 100ms baseline period of each trial.

### Analysis of MEG source data

In our source space analyses, we conducted a spatiotemporal cluster-based permutation test in both the right and left hemispheres to identify significant effects of our three factors (Combination, Locality, and Word order). We conducted this test as a 2 x 2 x 2 ANOVA in a time window spanning 100ms after target word onset through 450ms after target word onset (and corresponding to the onset of the following word). Because the target word was in a different location in the Adjective-Noun order and Noun-Adjective order stimuli, we shifted these conditions so that they were both time-locked to the onset of the target word. This simply means that in the Noun-Adjective condition, the baseline period was 2250ms before the onset of the target word, and in the Adjective-Noun condition, the baseline period was 2700ms before the onset of the target word. Though baseline correction can have a very strong effect on the amplitude of the word that immediately follows it, this difference is attenuated in later words in a sentence, and so by the time we measure the target word in the two Word order conditions, differences due to the baseline correction window were no longer clearly identifiable in the waveforms.

The goal of this analysis was to identify whether regions of the brain that typically show greater activation for words in Combinatory contexts relative to Non-combinatory contexts showed an interaction of Combination by Locality, and to identify regions that distinguished Locality or Word-order differences to guide questions for future research. We used a cluster p-value threshold of 0.025 in both the right and left hemispheres to minimize the risk of identifying false positive results, and we only consider clusters of at least 50ms in duration. We also apply Bonferroni correction (Rice, 1989) to the p-values that we report to account for the same test being performed independently in each hemisphere (Cabin & Mitchell, 2000). We report only the results that are below an alpha level of 0.05 after correction (corresponds to a p-value of 0.025 before correction).

### Decoding analysis

Our decoding analysis used a logistic regression classifier trained on sensor space MEG recordings to predict the what the trigger word was. For each comparison, we set up a 2-way contrast following (Fyshe et al., 2019). Our classifier was trained to distinguish each pair of two trigger words (e.g., distinguishing “blue” from “red”) separately. This means that, for each subject, we obtained accuracy scores for each of the six possible 2-way comparisons. Although it is also common to use vector representations of the target words with a linear classifier, the results end up nearly identical to our method given the small semantic prediction space generated by just four possible vocabulary items.

Following Honari-Jahromi et al. (2021), as part of our data processing pipeline, we first averaged together sets of three trials that shared a given trigger word (see Figure 3 for a diagram) to increase the signal-to-noise ratio (SNR). If a condition had a total number of trials that was not divisible by three, the excess trial(s) were excluded from that averaging block. In order to estimate the variance in decoding accuracy within each participant and increase the reliability of our accuracy measure, we did a Monte Carlo simulation and repeated this averaging with ten random permutations of the dataset, ensuring that trials that were dropped in one iteration could be included in others.

**Figure 3:**
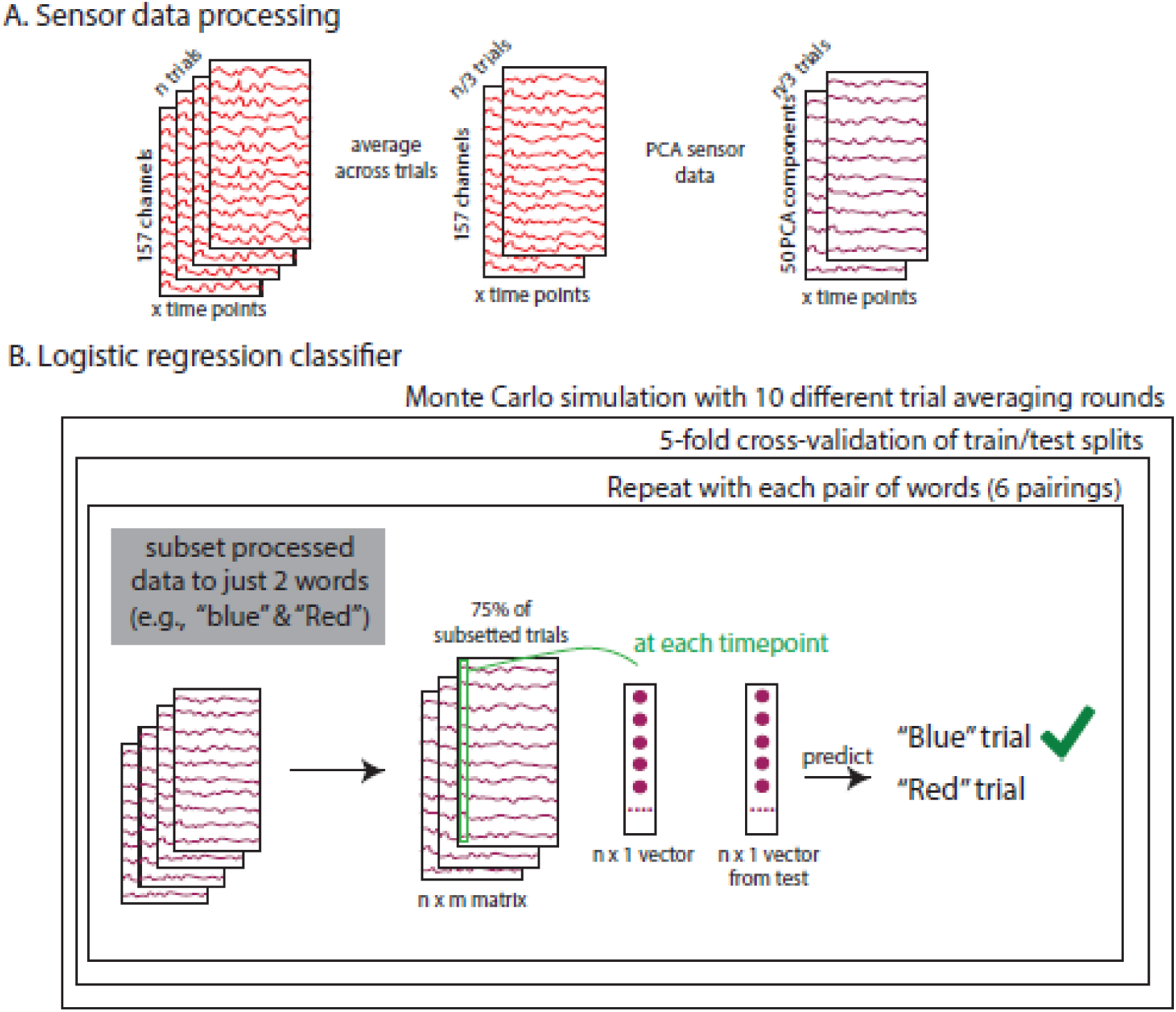
Diagram of the procedure used for the decoding analysis. (A) We take the filtered and cleaned sensor data and average together sets of 3 trials. We then use PCA to reduce the dimensionality of the data to the top 50 components. (B) Using separate logistic regression classifiers trained at each time point for each pair of two target words (6 total possible pairings), we measured classification accuracy. This process was repeated for 10 different random trial averages and used 5-way cross-validation in the train/test splits. In reporting results, we averaged together accuracy scores for each participant within each condition for each timepoint, and then calculate standard error across participants.

Following Dirani and Pylkkänen(2022), we then used PCA to reduce the dimensionality of the sensor data as a further data processing step. This step has also been shown to substantially improve the ability to decode complex information from sensor data (Grootswagers et al., 2017). We reduced the sensor data down to 50 dimensions, which accounted for at least 96.3% of the variance for each individual participant (mean 98.5%, median 98.7%). Finally, we scaled the data using Scikit-learn’s (Pedregosa et al., 2011) StandardScaler (see Guggenmos et al. (2018) for a discussion of the importance of scaling MEG data for decoding) and fit a logistic regression classifier with an L2 regularizer, using the SlidingEstimator function to get an accuracy score at each timepoint in the data using 5-fold cross validation of the train/test splits. We chose not to downsample along the time dimension or average time bins together as this reduces the effective sampling rate and can make classification more difficult in some cases (Rafidi, 2018).

## RESULTS

### Behavioral results

We analyzed both reaction time and accuracy results of participants while they were completing the main MEG task. We removed outlier reaction times that were greater than four seconds from this analysis (5.5% of all trials). Table 3 shows the mean reaction times and accuracy scores in different conditions. Each individual participant achieved an accuracy of at least 85% for the entire task, and most were well above that threshold, as the overall mean accuracy was 94%.

**Table 3:**
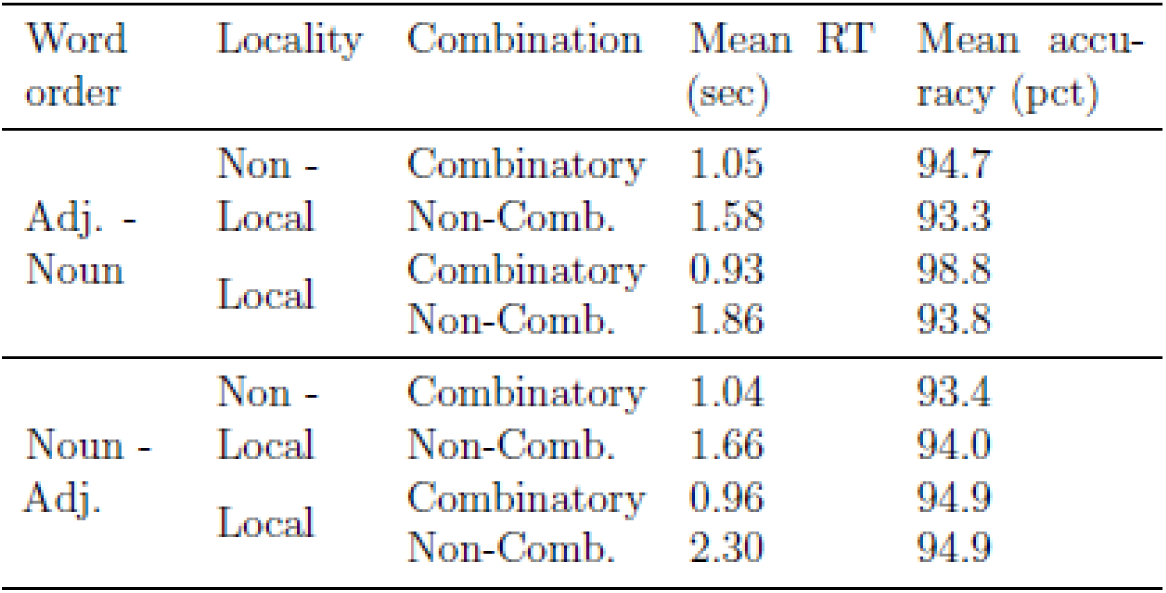
Table of behavioral results for all 22 participants during the main MEG task, broken down by each condition. There were significant differences in participants’ reaction times between Local and Non-local conditions and between Combinatory and Non-combinatory conditions, though accuracy remained high throughout. “Adj.” is short for “Adjective.”

Though the main questions of this study are only related to online processing measures, these results can provide insight into general differences in cognitive load across conditions for the stimuli that were used. There were no significant differences in participant accuracy between Locality and Combination conditions. In the reaction time data, a repeated measures ANOVA revealed a significant interaction of Locality by Combination, F(1, 22) = 173.8, p < 0.01. Follow-up pairwise comparisons showed that there were significant differences within both of those factors: the Non-combinatory items led to longer reaction times on the picture verification task compared to the Combinatory items (*F* (1, 22) = 515.21, *p* < 0.01), and the Local items had longer reaction times than the Non-Local items (*F* (1, 22) = 11.13, *p* < 0.01).

These reaction time differences may be the result of having to hold two or three different items in memory for the Non-combinatory items. There even seems to be a somewhat linear relation-ship between the number of nouns introduced in the sentence and the participants’ reaction times, as each of the Combinatory conditions introduced one object, the Non-Combinatory items in the Non-local condition introduced two objects, and the Non-Combinatory items in the Local controls were essentially a list condition that introduced three objects. It is difficult from just these results to determine if this is similar to the composition advantage effect (Rabagliati et al., 2017) because the task’s pictures for the sentences that introduced more than one object tended to also be more complex, on average, then the task pictures for the other conditions. So, picture complexity, rather than just sentence complexity, also may have led to longer reaction times.

These differences were very likely due to the increased cognitive load of having to use a list context in order to derive the Non-combinatory conditions. These differences do not necessarily reflect differences at the target word, as we assessed reaction times and accuracy only after the sentence was complete. Though some sentence conditions may have been more challenging than others for this task, there may not have been similar processing costs mid-sentence in the target regions for the MEG analysis. Additionally, when there were differences between Combinatory and Non-combinatory conditions, it was always the Non-combinatory items that appeared more difficult (lower accuracy; longer reaction times)—this is relevant because greater difficulty is often associated with greater activation in MEG recordings. Thus, any effect of greater difficulty in some conditions is likely to push the results *against* the deep composition hypothesis, which predicts an increase in activation for the Combinatory stimuli regardless of Locality. This difference was also the case with many of the previous studies of conceptual combination that have compared a list condition to a combinatory one (Bemis and Pylkkänen, 2011, i.a.). As with previous studies, it is also the case here that if the greater difficulty is obscuring our ability to measure conceptual combination effects, then this would likely not push results towards the locally-dependent composition hypothesis either, which predicts differences in conceptual combination in different Locality conditions, as the Non-combinatory items are also more difficult in the Local controls.

### Source space analyses: Spatiotemporal tests

#### Left Hemisphere effects

We observed a significant interaction of Combination by Word order in a left-hemisphere region that spans the anterior portion of the temporal lobe (*p_corr_* = 0.006; see Figure 4). This interaction was characterized by an increase in activation for the Combinatory condition relative to the Non-combinatory condition for the Adjective-Noun order; however, the direction of the effect was reversed in the Noun-Adjective order stimuli, and in this analysis we observed an increase in activation for the non-combinatory condition. In the pairwise comparisons, there were no significant effects that were solely attributable to the Locality factor. The follow-up pairwise comparisons showed that Combinatory and Non-combinatory conditions significantly differed from each other within each of the other 4 factorial conditions (all *p_corr_s* < 0.05, after correcting for 4 pairwise comparisons against the same null hypothesis).

**Figure 4:**
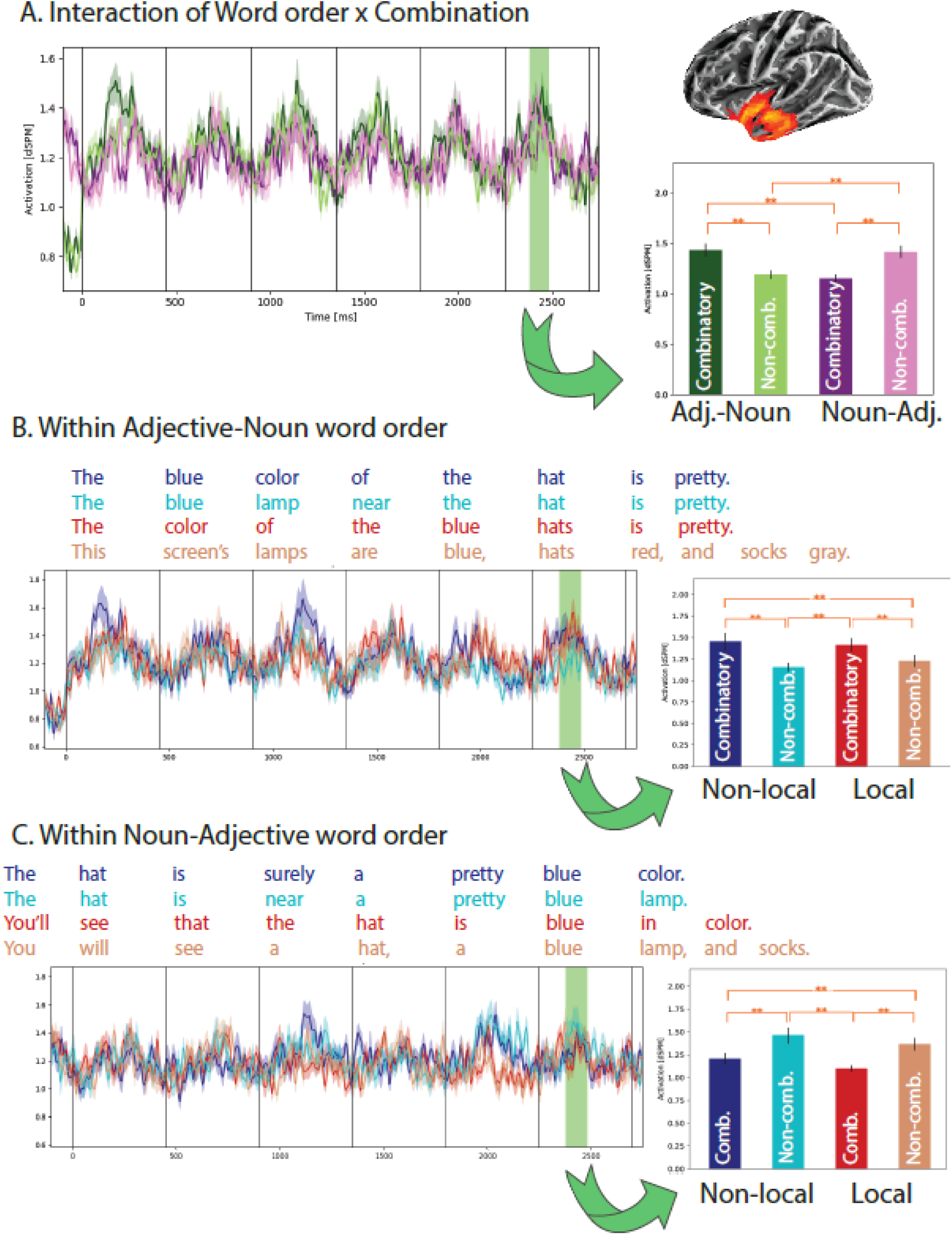
A. Primary results of the 2×2×2 spatiotemporal analysis showing greater activation for Combinatory stimuli in the Adjective-Noun word order condition relative to Non-combinatory stimuli, and the opposite effect in the Noun-Adjective Word order stimuli. B. Timecourse for the spatial cluster identified in part A with the same timecourse highlighted to show that this effect is not modulated by Locality within the Adjective-Noun word order condition, included primarily for visualization purposes. C. Timecourse for the spatial cluster identified in part A with the same timecourse highlighted to show that this effect is not modulated by Locality within the Noun-Adjective order stimuli either, included primarily for visualization purposes. All significance marking reflects pairwise comparisons that have been Bonferroni corrected to account for four tests against the same null hypothesis. **pcorr<0.01, *pcorr<0.05. ‘ indicates that the comparison was no longer significant after multiple comparisons correction.

For visualization purposes, we split the data in this identified cluster by the two Word order conditions to better visualize the effects of Combination and Locality. As these conditions do not interact in the main comparison, we only include the results of the pairwise comparisons to show that the observed effects of Combination were reliable, but we note that caution should be taken when interpreting these p-values, as the follow-up pairwise comparisons within each Word order condition was not licensed by the 2 x 2 x 2 test. As can be seen in the plots, however, the effects of Combination are not modulated by Locality within either Word order condition.

#### 2 x 2 x 2 Interaction

There was also a significant but complex three-way interaction of all the factors as well as an interaction of Word order by Locality (*p_corr_* = 0.03). This cluster covers part of the left vmPFC and IFG and extends from 250-335ms after the onset of the target word. The interactions in this cluster are not readily interpretable, and few pairwise comparisons survive multiple comparisons correction, so we include these results in more detail in Appendix D.

#### Significant main effects

We observed two separate main effects of Locality from the spatiotemporal analysis (100-190ms, *p_corr_*<0.001; 195-165ms, *p_corr_*=0.01). In both cases, the cluster localized to visual areas, including more occipital areas along the medial wall, and was characterized by an increase in activation for the Local items compared to the Non-local ones, possibly reflecting visual features of the word or relative changes compared to activation at the previous word. We did not observe any significant main effects of Combination or Word order in the left hemisphere.

#### Right Hemisphere effects

In the right hemisphere, we observed an analogous pattern of results to those in Figure 4, an interaction of Word order and Combination characterized by an increase in activation for the Combinatory condition in the Adjective-Noun order stimuli, but a decrease in activation in the Noun-Adjective order stimuli, relative to Non-combinatory controls. However, the cluster identified in the right hemisphere mostly localized to the medial wall, making the interpretation of the spatial extent of the cluster unclear (see plot in Appendix E). We also observed a significant main effect of Word order (*p_corr_* = 0.02; 195-265ms after target word onset, characterized by increased activation for the Adjective-Noun condition compared to Noun-Adjective) that mostly localized to the occipital region, and so this difference was due to visual features of the wordforms that varied systematically between the two conditions.

#### Decoding analysis

We plot the average accuracy of our decoding classifier in Figure 5. The shaded regions represent participant-wise standard error of the mean at each timepoint. We assessed significance using a temporal cluster permutation test within each word’s window for each Word order condition to analyze if the observed mean accuracy rates were significantly greater than the chance rate of 50% for a span of time. We computed a one-sample t-statistic, calculated by subtracting the chance rate of 0.5 from each sample timepoint to test if the value was significantly above 0. We plot all the significant clusters identified in Figure 5 as a shaded bar at the top of the plots with colors corresponding to the condition that was significantly above chance for that timespan. Regions that were significant below a p-value threshold of 0.05 are shaded in darker red and blue, and those time clusters that were only marginally significant are represented with the lighter-colored bars.

**Figure 5:**
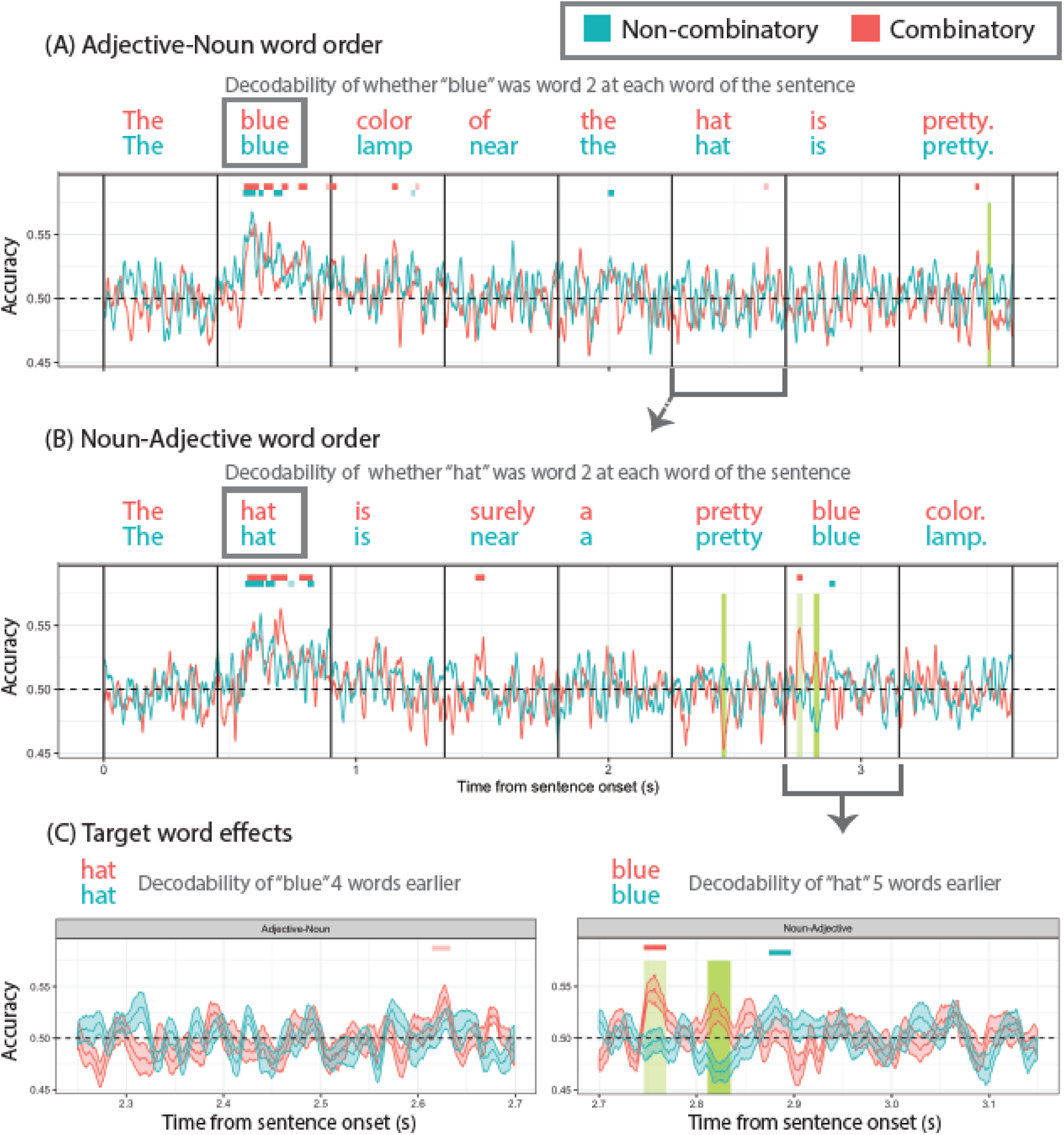
Results of the decoding analyses showing decodability of the trigger word at each point in the sentence. Chance accuracy is 50% and is represented by the horizontal black dotted line in all the plots. The black solid vertical lines are the onset of each word. The vertical green bars highlight the region where the decoding accuracy for the Combinatory and Non-combinatory conditions significantly differed from each other (darker green p < 0.05, lighter green represents a marginal trend at p < 0.075), and the lines on the top of the plot indicate when each condition showed significantly above-chance accuracy (darker colors are where p < 0.05, lighter colors represent a marginal trend at p < 0.075). (A) is just the Adjective-Noun condition, (B) is just the Noun-Adjective condition, and (C) zooms in on the decoding accuracy at the target word in each of the two Word order conditions. Error bars represent participant-wise standard error of the mean.

Unsurprisingly, we observed the highest classification accuracy when decoding for the trigger word as it was being presented, with all conditions showing around 55% decoding accuracy beginning around 100ms after the presentation of the trigger word. All four conditions showed a total of at least 110ms total of significantly above-chance decoding accuracy. However, these time windows were often slightly non-contiguous, which we assume is due to noise in the data or between-subject differences.

The primary region of analysis was the target-word window (“hat” in the Adjective-Noun stimuli; “blue” in the Noun-Adjective stimuli). In the Noun-Adjective condition, we observed above-chance decoding accuracy for the Combinatory condition 46-69ms after target word onset (*p* = 0.003). The Non-combinatory condition also showed significantly above-chance accuracy for a period of 174-197ms after target word onset (*p* = 0.021). In the Adjective-Noun word order, there was only a region where the Combinatory condition was marginally above chance from 364-383ms after target word onset (*p* = 0.07). The pattern of above-chance decoding accuracy for the Combinatory condition in the two Word order conditions represents very different latencies, and in neither case does the timing clearly line up with the times in which the Combinatory and Non-combinatory conditions differed in the source space analysis.

To determine whether accuracy significantly differed between the two conditions, we used a cluster permutation test with 10000 samples and a cluster *p*-value threshold of 0.05 to compare subjects’ data within each of the two Word order conditions separately, computing an *F* statistic for each timepoint. We found that decoding accuracy at the target word for the Combinatory condition was significantly higher than that of the Non-combinatory condition for two time periods in the Noun-Adjective word order condition at 111-135ms (*p* = 0.002) and 46-69ms (*p* = 0.054) after target word onset. This is represented in Figure 5 by the green vertical highlighting, with the lighter highlighting indicating the marginally significant effect. The accuracies in the Adjective-Noun condition did not significantly differ by Combination on the target word.

In the Adjective-Noun condition we also observed that trigger word “blue” could still be decoded on the word following its presentation for some brief but non-contiguous time periods. In the Combinatory condition, this was the case from 0-22ms (*p* = 0.026), 242-264ms (*p* = 0.006), and marginally 335-351ms (*p* = 0.059). In the Non-combinatory condition, there was only a brief period of marginal significance from 317-335ms after word 3 onset (*p* = 0.073). At the third word position, both “the blue color” and “the blue lamp” have identical structures, and we expect “blue” to compose with the noun that follows it in both cases, as both are combinatory at that point in the sentence (recall that condition naming refers to the composition at the target word).

Though not part of any of our hypotheses before beginning the experiment, we observed significantly above-chance decoding accuracy in the Combinatory Noun-Adjective condition for words like “surely” 121-159ms after word onset (*p* = 0.005, note that this also corresponds to an increase in LATL activity in the source space analysis). We also observed significantly above-chance decoding accuracy for the Combinatory Adjective-Noun condition on the fifth word (“the”) that extended from 196-221ms after word onset (*p* = 0.026). We did not observe any time windows with significantly above-chance decoding accuracy in the word 1 position, as is expected, since at that point in the sentence there was no information about what the trigger word (always the second word) will be.

## DISCUSSION

### LATL conceptual combination can operate non-locally

Overall, we found evidence in support of the *generalized composition hypothesis* for LATL conceptual combination. That is, the increase in LATL activity observed for words in Combinatory sequences compared to Non-combinatory ones reflects the conceptual combinations that are present in the composed conceptual representation of the sentence, irrespective of whether the composing concepts are adjacent or merge with each other syntactically. This suggest that conceptual combination in the course of language processing is, to some extent, its own processing tier, not faithfully following the syntax.

Our decoding analysis further showed that in Combinatory contexts, but not Non-combinatory contexts, it is possible to decode from the neural signal which word is being composed even when that word occurred several words previous, and even when there have been intervening content words. Crucially, this study provides the first direct evidence for reactivation of a specific lexical item for the purpose of composition, and it does so in a construction that does not require any specific gap construction, indicating that the retrieval is not necessarily syntactic, but rather a part of a meaning building operation.

Thus, we have the broad generalization that, when measuring on a target noun, we observe an increase in LATL activity for composition. And when we track the representation of a noun throughout a sentence, we find that that noun is reactivated at the point at which it needs to compose with a later adjective. Taken together, these findings indicate the parser selectively reactivates lexical representations, that this reactivation can be dependent on whether that representation is needed for composition, and that the LATL’s role in language processing is a generalized combiner. A puzzle that emerged from this study pertains to the effect of word order, which flipped the directionality of the composition effect. We discuss the reversal in the third section below, though ultimately, further studies are required to understand it.

### Implications for conceptual combination

This study found evidence that conceptual combination relies on meaning-based cues that indicate which concepts need to be combined. Previous work had shown that the semantic and conceptual properties of words could attenuate effects related to conceptual combination, indicating an importance of the parser’s linguistic knowledge in conceptual combination (refs). This study took a different approach to showing how conceptual combination interacts with linguistic knowledge by relying on word-level cues to indicate whether two words in separate syntactic phrases should be combined. We showed that, at least in the Adjective-Noun configurations used, an adjective from a separate grammatical phrase than the noun it modifies will still lead to an increase in activation in the LATL, relative to that same adjective and noun in two phrases that form a non-combinatory context. This shows that the LATL’s role in conceptual combination is more general than was previously known.

Even though the brain is sensitive to local phrase structure in composition (Mollica et al., 2020), we have shown that two lexical items that are part of separate syntactic phrases can engage the LATL in a very similar way to how those two words would if they were part of the same phrase. This extends previous findings from Parrish and Pylkkänen (2022) which showed that the LATL is engaged in conceptual combination for two words that do not syntactically merge (but are otherwise part of the same DP structure). This study furthers our understanding of how conceptual combination fits in to a larger parsing model as well as narrowing down the possibilities of how the mechanism behind conceptual combination operates. Conceptual combination in the LATL not only does not depend on syntactic merge, it does not depend on the composing words even being part of the same constituent. We suggest that the relationship between syntactic processing and conceptual combination is one in which sentential information makes different words available for composition (e.g., the possessive structure of “the blue color of …” allows the possessum “blue color” to remain available for composition, but the locative structure of “the blue hat near…” does not), and the LATL is engaged when composing any reasonably-available words.

The findings of the decoding analysis further support this explanation of the relationship between structure and conceptual combination. If, in fact, meaning-based cues in a sentence alter what words are available for composition at different time points, then this suggests that the mechanism responsible for conceptual combination does not need to distinguish how a given lexical representation became available to it. Indeed, in the source space analysis, we observed similar increases in activation for nouns in combinatory contexts relative to non-combinatory contexts that was identical for local composition (where the adjective and noun were linearly adjacent) and long-distance composition, where three words intervened between the adjective and noun. And the decoding analysis provided evidence that the different sentence types affected which words were decodable from the neural signal, with earlier nouns reactivating very early in processing in a way that is linearly decodable specifically when they are needed for composition.

On a high level, we suggest the mechanism behind conceptual combination takes part in the process of selectively composing words. This does not, of course, mean that the LATL is directly involved in tracking which representations should remain available for composition at different points in a sentence. It means that the observed relationship between LATL activation and composition participates in the broader parsing process in which this takes place, as opposed to the effect being solely present to build minimal phrases. Although we cannot make a causal claim about the role of the LATL in meaning composition, these findings suggest that the region is dynamically acting on input made available to it both from the internal parsing/memory state, and from external linguistic stimuli. It remains an open question whether conceptual combination fully ignores incoming input if it cannot yet be integrated into the conceptual structure being built, as we have only measured a relative difference in activation patterns between combinatory and non-combinatory contexts, though Parrish and Pylkkänen’s (2022) and Li and Pylkkänen’s (2020) findings strongly suggest that the incoming input is not ignored, even when the resulting composition would not result in a grammatically licit phrase.

This study is one of only a few to use predicate structures as a direct combinatory comparison to adjective-noun sequences. Our finding that Non-combinatory stimuli led to an increase in LATL activation for the Noun-Adjective order stimuli is surprising in light previous. Matar et al. (2021) used a minimal phrase paradigm in Arabic to compare predicate structures meaning “the car is red” to noun phrase structures meaning “the/a car is red,” and they found that both types of phrases similarly activated the LATL, in contrast to this study’s findings in the Noun-Adjective ordered stimuli. The stimuli that we used in this study were matched for their conceptual representations, but not necessarily matched for other aspects of the sentence that might affect processing. So, it is possible that the Noun-Adjective items were meaningfully different from the Adjective-Noun items, especially as the target word in the Noun-Adjective order stimuli strongly predicted that a noun (with which the target word had to compose) would follow, but this was not the case in the Adjective-Noun order stimuli. We expand on the questions raised by our Word order contrast in the following section.

### Word order differences

For generality, our manipulation contained not only the adjective-noun order, but also the noun-adjective order, both in local and long-distance configurations. Within both orders, composition affected LATL activity in the same way for local and long-distance stimuli, but surprisingly, the noun-adj order elicited an activity *decrease* in the presence of composition. Though with the present data, we are not able to determine why this reversal happened, there are several future directions that can be explored in subsequent studies. First, it is possible that structural or lexical prediction played a role. In the Noun-Adjective condition, it was clear from the sentence frame that the adjective could not yet fully compose with the earlier noun. We originally selected a set of several color-defining words (“color,” “hue,” “tint,” “shade”) to decrease the overall predictability of each word. However, this may have had the unintended effect of causing a delay in conceptual combination if the parser waited to find out if the hat was a lovely blue hue as opposed to a lovely blue color. Such an explanation would be at least partially in line with Parrish and Pylkkänen’s (2022) findings, where they found that syntactic cues indicating a structure was not yet complete led to an overall decrease in LATL activity in a time window compatible with conceptual combination. Though these syntactically incomplete stimuli still showed greater activation than the Non-combinatory controls, the lowered activation means that perhaps the stimuli used in this experiment were not the cleanest contrast. We suggest that perhaps the parser did not finish composing the phrase because the DP wasn’t finished yet in the Noun-Adjective condition, and that this cue to not compose is different from the way a potentially analogous cue would act in the presence of an incremental mismatch in syntactic number features of the type discussed in by Parrish & Pylkkänen (2022).

As another possibility, it may be that the structural frames that we used induced an additional processing load. As our Local condition used embedded phrases (the proposition “the hat is blue” was embedded under “you’ll see that”), it may be that embedded propositions contribute differently to updating conceptual representations in analogous non-embedded phrases. Such a finding would be surprising in light of our earlier conclusion that LATL conceptual combination is a very general component of language processing. This possibility would also fail to explain why the Non-local Combinatory condition also showed lower activation than the Non-combinatory condition, but as embedding structures have not been studied with respect to conceptual combination, we point this out as a potential factor.

We also consider that the use of a list condition for the Local controls had the effect of increasing activation more generally, compared to the Non-local Combinatory items which did not use a list. For this to be the case, it would also have to be true that the way we created the lists in the two Local Non-combinatory conditions were meaningfully different between the two Word order conditions. This is not out of the question, though, as the Adjective-Noun word order controls used a gapping construction (“This screen’s lamps are blue, hats red, and socks gray”), while the Noun-Adjective order ones needed to include a function word between the trigger word and the target word (“You will see a hat, a blue lamp and toys”). We noted in the behavioral results that the Local controls in the Noun-Adjective case were associated with the longest reaction times, indicating that the processing load for these sentences was likely overall much higher than in any of the other conditions, including the Adjective-Noun word order’s Local Non-combinatory condition. Within the Non-local Noun-Adjective condition, the Non-combinatory items also led to longer reaction times on average compared to the Combinatory items, but this difference was not necessarily any more pronounced than it was in the Adjective-Noun condition.

If this explanation is correct, then the high cognitive load of the list-type stimuli in the Noun-Adjective controls increased activation in such a way that the composition effects were overshadowed by this much larger effect. As part of an additional post-hoc analysis, we ran a spatiotemporal test in the left hemisphere contrasting the Combination and Locality conditions just within the Noun-Adjective order stimuli to determine if the effect of Combination localized more to the LIFG or other regions associated with memory or task load effects. In this analysis, we observed a main effect of Combination in which the Non-combinatory items showed greater activation than the Combinatory ones (as expected from the main source space analyses). While the spatial extent of this main effect extended slightly into the LIFG, it most strongly localized around the middle temporal lobe, and does not clearly support an explanation based on general cognitive load differences accounting for the observed effects.

Finally, it is always possible that predication does, in fact, dissociate from Adjective-Noun conceptual combination in some important ways. While within each Word order, the structures are very well matched, there is at least one very important difference between them. In the Adjective-Noun order condition, the two composing words are part of the subject of the phrase (though, a complex subject with multiple phrases). However, in the Noun-Adjective condition, only the trigger word is part of the subject, and the target word is part of the predicate phrase. However, one previous study using minimal subject-predicate structures did find an increase in LATL activity for the predicates in combinatory contexts relative to a list condition (Phillips and Pylkkänen, 2021), though they were measuring activity on the verb of the two-word sentences.

### Modification, predication, and possession

As the full sentential stimuli are complex and involve many different types of modification, it is worth additionally considering what effects these may have had, and we conduct an additional exploratory analysis to characterize these potential differences. In the Non-local Adjective-Noun order stimuli, there are two potentially very different types of modification at the third word in the sentence. In the Combinatory condition, we used “blue color” as opposed to “blue lamp” in the Non-combinatory condition to indicate that the concept of “blue” would still be available to compose with a later noun. However, this brought about an additional difference, as in “blue lamp,” the color adjective is used intersectively (the thing is both a lamp and is blue), whereas in “blue color” the color adjective is not clearly being used intersectively, at least not in an informative way (being both a color and blue does not add to the meaning of the phrase). Ziegler and Pylkkänen (2016) have shown that early LATL activity corresponds to intersective modification (“red boat”), while context-dependent scalar modification (“large boat”) corresponds to a later and slightly more posterior effect. In our case, the “blue” of “blue color” is not context-dependent, but rather part of a subsective relationship that may be different from intersection.

However, our post-hoc analysis found that the potentially non-intersective examples showed greater activation than the more straightforwardly intersective ones. We conducted an additional spatiotemporal test using a one-way ANOVA in the left hemisphere to compare the Combinatory and Non-combinatory conditions of just the Non-local Adjective-Noun order stimuli in word 3 position (comparing on “color” vs. “lamp”). The post-hoc test revealed that there was a significant difference between the two Combination conditions characterized by an increase in activation for “color” compared to “lamp” in much of the left temporal lobe and extending into the vmPFC 235-405ms after word onset (*p*=0.001), but we note that the design of this study did not intend to take this difference into account, and it’s possible that a confounding factor such as lexical frequency is driving the increased activation. The words that could be slotted in for the “color” word (“hue”, “tone”, “shade”, “color”) have, on average, a lower corpus frequency compared to the list of target nouns (“lamp”, “book”, “chair”, “hat”); the average frequency of the target nouns in COCA (Davies, 2010) was 2.5 times higher than that of the color words. The post-hoc analysis suggests that the LATL may be sensitive to differences between subsective or redundant modification and intersective modification, but a more carefully controlled study is needed. Further, this increase in activation on “color” is especially surprising in light of Blanco-Elorrieta and Pylkkänen (2016) findings that numeral modification (e.g., “two boats”) does not lead to an increase in LATL activation, as they argue that the reason for this is due to the numeral not adding to the conceptual content of the noun. In our study, it is also the case that “blue” does not clearly add to the conceptual structure of “color,” though it does substantially narrow down the conceptual content of “color.”

In addition to the different types of modification, the “color of the hat” constructions imply a kind of possessor relation. Previously, Westerlund et al. (2015) had used a minimal phrase possessive structure (e.g., “Tarzan’s vine”) to investigate argument saturation effects in LATL composition. Though they aggregated the results of possessive structures with other phrase types that indicated argument saturation (verb + noun and preposition + noun) in their analyses, they found that the argument saturation cases had a similar response profile in the LATL as their modification examples which included adjective + noun modification.

To return to the issue of modification vs. predication, Flick and Pylkkänen (2018) used a particular English construction that allows post-nominal adjectival modifiers (“the mountain tall enough for a strenuous hike”), but they did not contrast combinatory and non-combinatory contexts—rather they contrasted predication vs. modification by placing the phrase in a question (to create the predicative structure for “are many mountains tall enough…”) vs. an existential construction (to create the modification structure for “there are many mountains tall enough…”).

They found no difference between these conditions in the LATL on the adjective “tall,” but they did find that there was an increase in activation on the following word, “enough” in the modification examples relative to the predication examples, such that the modification examples showed significantly greater activation, suggesting additional differences between the two composition types. Ultimately, we conclude that differences between modification and predication for conceptual combination remains an open one, and future work teasing apart the contributions of each for LATL conceptual combination could be a valuable contribution to both our theoretical and psychological understanding of conceptual combination.

### Changes in decodability of words throughout a sentence

The paradigm used in this study to measure when different words in a sentence are active is one that, to our knowledge, has not yet been widely used. In addition to the main findings of this study, we observed effects as part of an exploratory analysis that merit follow up work. Notably, we observed that in the Noun-Adjective order stimuli, the earlier noun was decodable with above chance accuracy at the presentation of the adverb (“surely” in the example sentence). The words that filled this slot (“surely,” “clearly,” “definitely,” “indeed,” and “certainly”) were all high adverbs—that is, semantically they take scope over the entire sentence. We did not consider any hypotheses about this difference before running the study, so all discussion related to the findings in this region should be considered a post-hoc explanation, and we simply suggest a reasonable possible explanation for these findings. We propose that the reactivation of the earlier word in these examples may be similar to reactivation at sentence wrap-up that has been reported by Rafidi (2018). As high adverbs need to take scope over the semantics of the entire proposition, it may be that these words trigger reactivation because they scope over the previous words.

Some of the other observed effects may shed light on follow-up questions about the properties of already-composed structures. Specifically, we consider whether the properties associated with “blue” were still identifiable when the relevant concept is the already-composed phrase “blue lamp.” In the sentence “the blue lamp near this hat is pretty,” we assume that composition between “blue lamp” and “pretty” requires conceptual combination, and thus “blue lamp” should be re-activated at the final word in the sentence for composition. The question is then whether there is any sense in which the features of “blue” remain active after having composed with “lamp,” such that the example sentence will be more similar at “pretty” to a sentence like “the blue chair near this hat is pretty” compared to “the green chair near this hat is pretty.” Though there were two significant clusters identified at the presentation of “pretty” in these sentences (see Figure 5), the results were inconclusive. We would have predicted above chance accuracy in the Combinatory condition as “pretty” modifies “blue color,” and we do see a brief period of above-chance decodability for the Combinatory condition, but it occurs much later than the reactivation effect observed in the Noun-Adjective condition (though it is temporally similar to the marginally-significant re-activation at the target word in the Adjective-Noun condition). It may be that this is because “pretty” does not directly modify “blue,” but rather is predicated of “blue color,” the temporal dynamics of reactivation are different from the relationship observed in the Noun-Adjective condition.

We tested whether a classifier trained on the Non-combinatory condition would show above-chance accuracy, as that is the condition in which “blue” needed to previously conceptually compose with an object (it is still an open question whether “blue color” constitutes conceptual combination). We did observe a period where the Non-combinatory condition had significantly higher decoding accuracy compared to the Combinatory condition at the last word (“pretty”), but the Non-combinatory condition never had significantly above-chance decoding accuracy, making interpretation of this effect more difficult. It would not be justified to claim that the earlier concept of “blue” was active at the final word of the sentence in the Non-combinatory condition, at least not in a way that we were able to measure. However, the fact that there was a reliable difference between conditions means that this question is worth following up on in future work, particularly any work that can use a more robust contrast to decode for, as it may simply be that in a full sentence context, the representations for “blue” as opposed to “green” were not sufficiently distinct to answer these questions.

### Directions for future work

This study both advanced our understanding of the generalizability of LATL conceptual combination and raised new questions. Future work is needed to better understand compositional differences between adjective-noun composition and predicative modification, as the difference that we observed in this study was surprising, especially in the Local control condition. It will be important in future studies to better control for effects of structural or next-word prediction, a possible confound from this study. If adjectives in predicative relationships to a noun are different from adjectives as prenominal modifiers, then this would provide an important challenge to our account of conceptual combination as a very general combinatory operation.

In our decoding analysis, though we observed lexical re-activation in our combinatory condition, decoding accuracy was ultimately low throughout the experiment and never reliably reached above 60%, even in cases where we were decoding which word was being presented at that time. Thus, a more robust contrast may be needed for probing the questions introduced in this study. We suggest that future studies make use of a stronger contrast, such as contrasting two semantic categories of nouns (e.g., Dirani and Pylkkänen (2022) contrast tools and animals) or high vs. low specificity of adjectives. It may be that the representations for “blue” and “black” were too similar for us to distinguish them reliably, but it is reasonable to think that a contrast like “blue” vs. “striped” might be easier to detect at the target word.

This study also provided a kind of proof of concept for using decoding analyses to probe syntactically-cued lexical reactivation. This kind of a paradigm has exciting implications for our understanding of other types of long-distance dependencies. As previously mentioned, much of the work on long-distance dependencies has focused on filler-gap constructions, and many questions remain about the nature of what kind of a representation gets reactivated. If this kind of a paradigm can reliably allow us to track with high temporal resolution which representations are active when, this design could help address long-standing questions about the kinds of syntactic cues that trigger maintenance and retrieval operations. And tracking the activation of individual lexical items could open the door to testing general processing accounts of syntactic islands.

## CONCLUSION

This work addressed the nature of conceptual combination during language processing, testing whether instances of conceptual combination are observed over a long distance for concepts that do not syntactically combine with each other. Our hypotheses pertained to the left anterior temporal lobe, gives its well-documented role in conceptual combination, although our analysis did cover the whole brain. While word order had a surprising effect, the left anterior temporal lobe did show a clear sensitivity to conceptual combination across word orders with no interaction with locality, that is, the same effect was observed for both local and long-distance conceptual combination. Thus, our findings show neural evidence for the existence for long-distance conceptual combination and implicate the left anterior temporal lobe as the site where this process is implemented. Further, our decoding analysis showed that earlier nouns reactivate for a brief period in combinatory contexts, but not non-combinatory ones, suggesting that long-distance conceptual combination may operate via concept reactivation.

## Acknowledgements

This work is funded by NSF grant BCS – 2140741 and NYU Abu Dhabi Institute Grant G1001.

## Appendix A. Stimuli creation details

**Table 4:** Norming results from the experiment. The phrases represent the set of sentential phrases that fit a certain frame. For example, “book above x” includes sentences that state a book is above a lamp, chair, key, and hat. This relation can occur in phrases of the form “The book above the lamp is…” or of the form “The book is above a lamp.” The values in the “Percent > 4” column represent the percentage of the time that the average participant rating for a given sentence in that configuration was strictly above 4 on a 7-point rating scale.

| Phrase<br>(Noun position) | Percent > 4 | Phrase<br>(Position noun) | Percent > 4 |
| --- | --- | --- | --- |
| "book above x" | 100 | "x above book" | 75 |
| "book below x" | 25 | "x below book" | 44 |
| "book near x" | 100 | "x near book" | 74 |
| "book on x" | 86 | "x on book" | 50 |
| "book under x" | 60 | "x under book" | 100 |
| "chair above x" | 45 | "x above chair" | 60 |
| "chair below x" | 50 | "x below chair" | 50 |
| "chair near x" | 100 | "x near chair" | 100 |
| "chair on x" | 36 | "x on chair" | 75 |
| "chair under x" | 55 | "x under chair" | 33 |
| "hat above x" | 56 | "x above hat" | 75 |
| "hat below x" | 67 | "x below hat" | 60 |
| "hat near x" | 89 | "x near hat" | 100 |
| "hat on x" | 60 | "x on hat" | 60 |
| "hat under x" | 60 | "x under hat" | 57 |
| "lamp above x" | 86 | "x above lamp" | 67 |
| "lamp below x" | 44 | "x below lamp" | 62 |
| "lamp near x" | 100 | "x near lamp" | 100 |
| "lamp on x" | 50 | "x on lamp" | 55 |
| "lamp under x" | 50 | "x under lamp" | 25 |

**Table 5:** All vocabulary inserted into templates to create stimuli.

| Word type | Vocabulary items |
| --- | --- |
| TARGET NOUN | lamp, hat, book, chair |
| TARGET COLOR | blue, red, green, black |
| POSITION WORDS | near, above, below, on, under |
| VERBS OF SEEING | see, notice, find |
| ADVERBS | surely, clearly, definitely, indeed, certainly |
| ADJECTIVES | pretty, lovely, nice, neat, stylish |
| INTENSIFIERS | very, really, quite, rather |
| COLOR DESCRIPTORS | color, hue, tint, shade, tone |
| SCREENS | screen, display, image, picture |
| NON-TARGET COLORS | pink, white, brown, purple |
| NON-TARGET NOUNS | toys, boxes, bikes, scarves, socks, pens, bags |

**Table 6:**
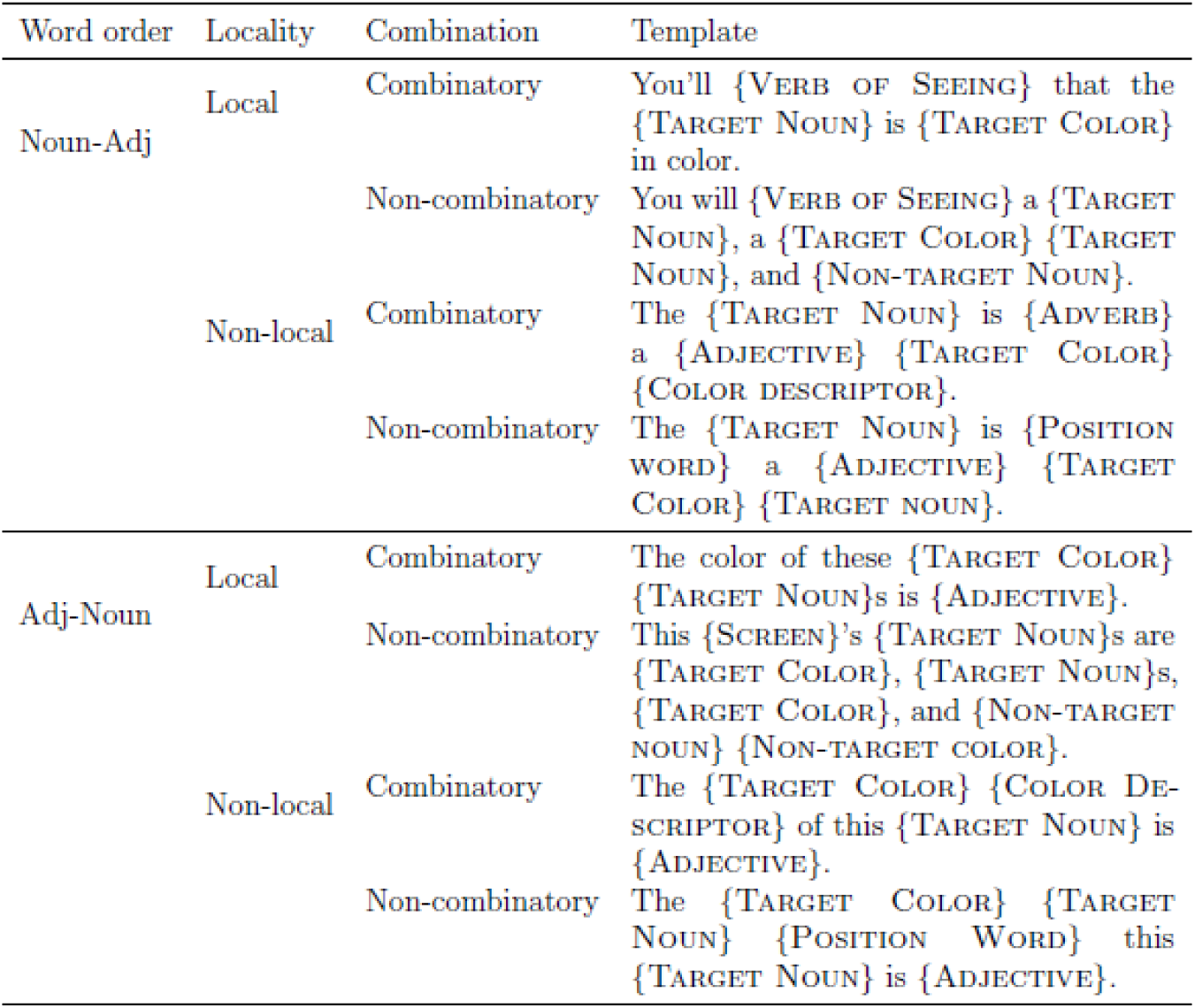
All eight templates used in creating stimuli. The Target Noun and Target Adjective were always balanced across stimulus sets. All other vocabulary items were slotted in to increase lexical diversity and decrease the repetitiveness of the task, and thus they were not perfectly controlled for.

## Appendix B. Stimuli norming

### Instructions and Interface

The two examples had the correct radio button already highlighted, and they were followed by an explanation of why that selection is correct. Additionally, participants were warned that there would be attention check items throughout (described in the main text), and they were given explicit instructions about how to treat the subset of attention check items that included a number, such as “Select number five to show that you are still paying attention.” We added these explicit instructions because we determined which work we would accept or reject (and thus which work we would pay participants for) based on whether they passed these instructed response attention check items, and we wanted to make sure that it was completely clear to anyone making an honest effort at the task what was expected of them.

Each of the questions were presented as short sentences that the participant rated using the 1-7 likert scale. An example of this set-up can be seen in Figure 7, which shows a screenshot of the norming task. The first four of 21 items are shown; item (2) is an example of a filler item that was semantically mal-formed, and item (4) is an example of a filler that was an instructed-response item. The order of presentation of the items in was randomized. The survey was written as a plain html form and presented natively in the MTurk interface, so that participants did not have to leave the MTurk site to participate in our norming study.

**Figure 6:**
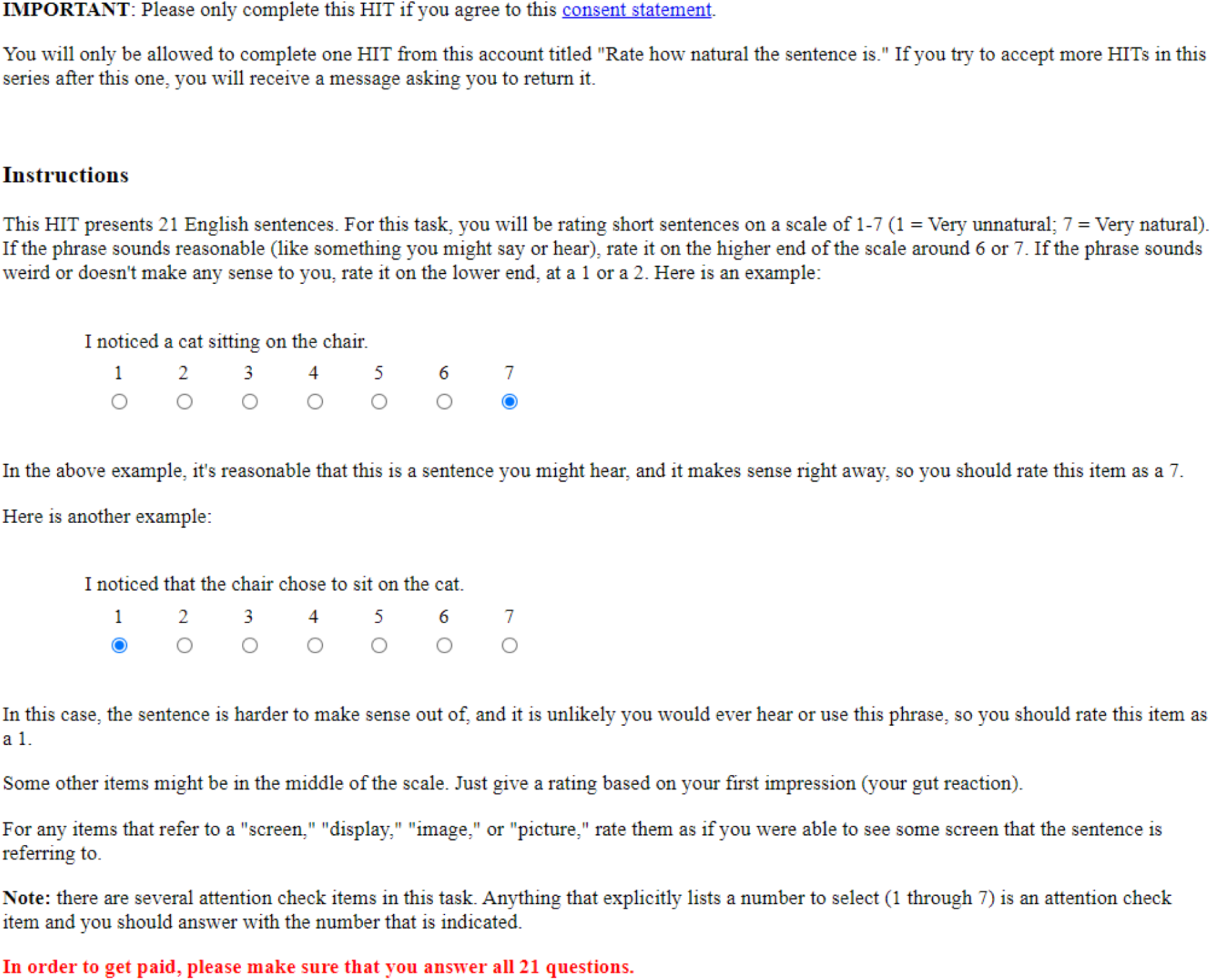
Example screenshot from the MTurk rating task. This figure shows the instructions that all participants saw before beginning the experiment.

**Figure 7:**
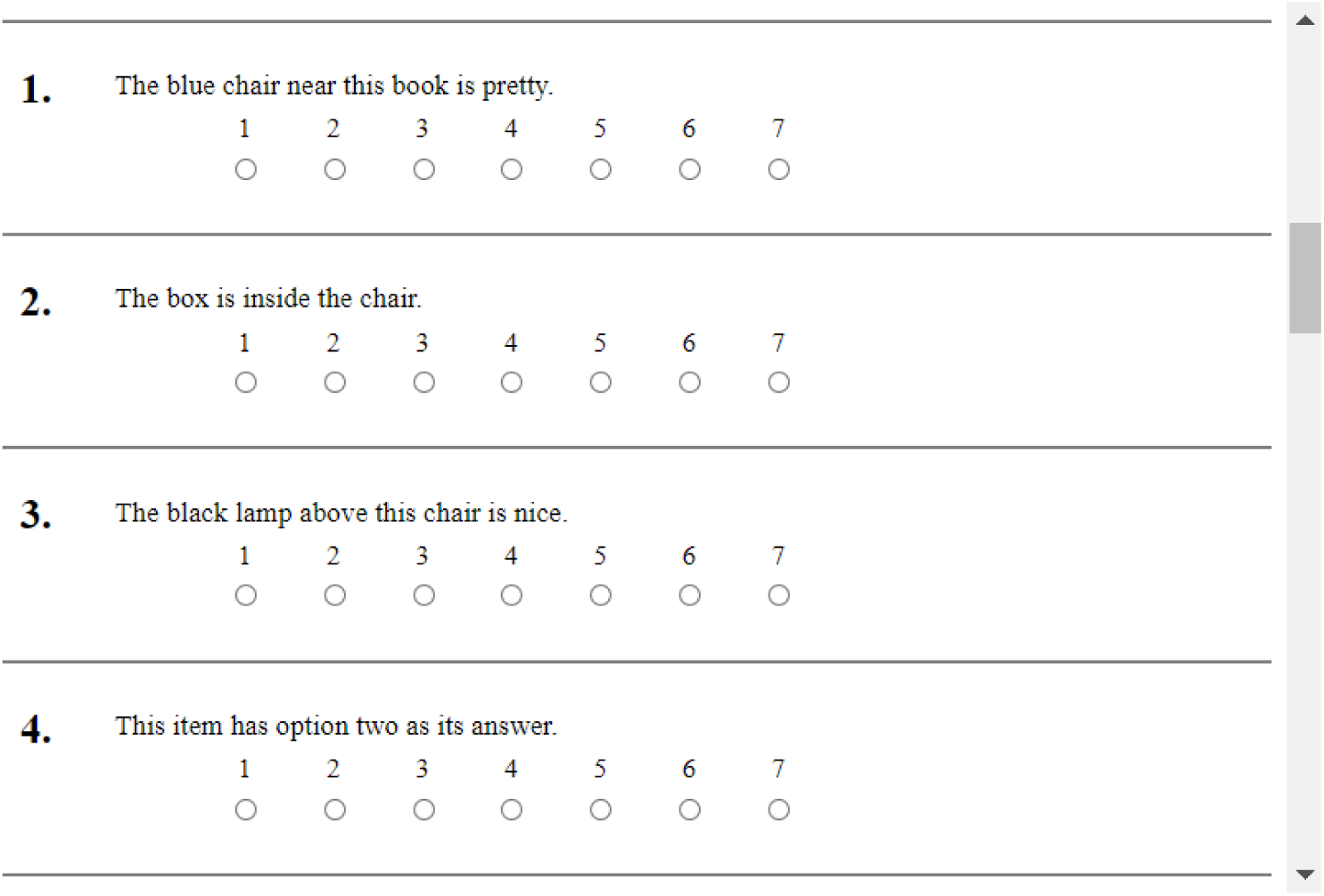
Example screenshot from the MTurk rating task. This figure shows the first four items that were shown to one participant.

### Norming results

Participants rated all possible combinations of target nouns and positions that those nouns could occur in. In order to assess which target nouns are natural in which configurations, we broke down the results into which nouns were rated above a 4 (on the 7-point scale of naturalness) when presented in either the first slot (noun is position + x) or second slot (x is position noun) relative to the position word. The results in Table 4 shows the results of this breakdown, presented as the percentage of times that a word in each position was rated, on average, as acceptable in that position. We considered ratings strictly above 4 to be “acceptable” for the purpose of this study. According to these results, for example, it was relatively unnatural to state that a chair is on another target noun (36% acceptable), but relatively natural to say that another target noun is on a chair (75% acceptable).

We used these results to exclude the most unnatural combinations, with the assumption that the unnatural ones had much lower transitional probabilities given our small vocabulary, and increasing the predictability of any words, but especially the target nouns, could make the composition effects we were testing for less likely to occur, presenting a potential confound in interpreting the results, as our Non-combinatory condition in both Word order conditions relied upon the position word. Therefore, we only included position relationships that were rated as acceptable (average ratings were above 4) more than half the time, as this indicated that there were multiple possible nouns that could satisfy a given relation. All other relations were excluded when we used the templating script to create our experimental stimuli.

### Attention checks

As described in the main text, we included 7 attention check fillers for each participant who took part in the norming study. To ensure that participants were not able to share answers between each other on online forums, we used a randomly assigned set of six such items, ensuring that three of the attention checks were instructed response items (one from each answer range) and three were gold-labeled items (one from each category).

We show the response rates of 272 of the original 279 participants who completed the task. We removed data from the 7 participants whose work we rejected for failing to pass at least 2 of the instructed response items, as we assume that these participants were not reading the sentences and simply answering at random. We show the average results across all participants in Tables 9 and 10 to show that the attention check items were appropriately discerning which participants were reading the sentences closely enough to be included in the analysis.

**Table 9:** Table of all instructed response items used in the norming study. Each participant saw one of the items from each of the three answer ranges.

| Text | Ans.<br>Range | Accept.<br>range | Mean<br>resp. | Acc.<br>rate |
| --- | --- | --- | --- | --- |
| The key to completing this attention check is selecting six. | High | 6 | 5.98 | 97.56 |
| This attention check item has option six as its answer. | High | 6 | 6 | 100.0 |
| The option to select is seven on this item. | High | 7 | 7 | 100.0 |
| They say that you must select seven to show you are reading. | High | 7 | 6.99 | 98.70 |
| The option to get this item right is option seven. | High | 7 | 6.89 | 95.74 |
| The key to getting this attention check right is selecting two. | Low | 2 | 2.11 | 96.30 |
| This item has option two as its answer. | Low | 2 | 2.17 | 96.04 |
| The option to select is one on this attention check. | Low | 1 | 1 | 100.0 |
| They say this is an attention check, select one here. | Low | 1 | 1.03 | 98.33 |
| The option to get this item right is option one. | Low | 1 | 1 | 100.0 |
| To show that you are paying attention select three. | Mid | 3 | 3.03 | 98.33 |
| To pass this attention check select four. | Mid | 4 | 4.13 | 95.83 |
| To show that you are paying attention select five. | Mid | 5 | 5 | 100.0 |
| To pass this attention check select five. | Mid | 5 | 5 | 100.0 |
| To complete this attention check select item four. | Mid | 4 | 4 | 100.0 |

**Table 10:** Table of all gold-labeled items used in the norming study. Each participant saw one question from each of the three categories. Sentences in the “Good” category were written to be acceptable, those in the “Sem.” category were written to be unacceptable for semantic reasons, and those in the “Pos.” category were written to be unacceptable due to the position word.

| Text | Category | Accept.<br>range | Mean<br>resp. | Acc.<br>rate |
| --- | --- | --- | --- | --- |
| There is one dog barking in the garden. | Good | 6-7 | 6.42 | 83.33 |
| I can see that the toy is on the floor. | Good | 6-7 | 6.03 | 66.67 |
| Those purple toys are laying on the floor. | Good | 6-7 | 6.62 | 89.66 |
| The box is definitely still in good condition. | Good | 6-7 | 5.68 | 69.05 |
| Those dogs have black spots and those cats do not. | Good | 6-7 | 6.78 | 96.30 |
| There are no dogs outside and they are all barking. | Sem. | 1-2 | 2.33 | 71.43 |
| Those dogs have black spots but those same dogs do not. | Sem. | 1-2 | 2.32 | 65.85 |
| No dogs are outside and they are all playing. | Sem. | 1-2 | 2.82 | 65.00 |
| I can see that the bag is open and the bag is closed. | Sem. | 1-2 | 2.25 | 72.28 |
| I can see that the floor is on the toy. | Pos. | 1-2 | 1.49 | 87.95 |
| The floor is laying on those purple toys. | Pos. | 1-2 | 1.78 | 83.3 |
| The box is quite still in a green condition. | Pos. | 1-2 | 1.83 | 73.9 |
| The box is inside the chair. | Pos. | 1-2 | 2.4 | 69.41 |
| The picture is hung up on the floor. | Pos. | 1-2 | 1.92 | 81.94 |

## Appendix C. Single-word picture-verification task

In order to increase the number of trials we could use in the decoding analysis, we introduced a short task that participants completed before the main full-sentence picture-verification task. In this task, we measured MEG activity while participants read a word presented on the screen. Following the word, the participant saw a picture, and their task was to answer, via button press, “yes” if the picture matched the word, and “no” otherwise. The stimulus presentation timing was identical to that of the main task to ensure that the single-word task would be maximally generalizable to the main task. Participants each completed 240 trials, which took approximately six minutes. Figure 11 shows examples of trials for both the Noun and Adjective stimuli, though in the task these were not separated into different sections. Any time participants were presented with a Noun, the picture was one of the four possible objects, and any time they were presented with a color word, the picture that followed was a blob of one of the four possible colors. Half of the trials had a correct answer of “yes,” and the order of presentation of the trials was fully randomized for each participant.

**Figure 8:**
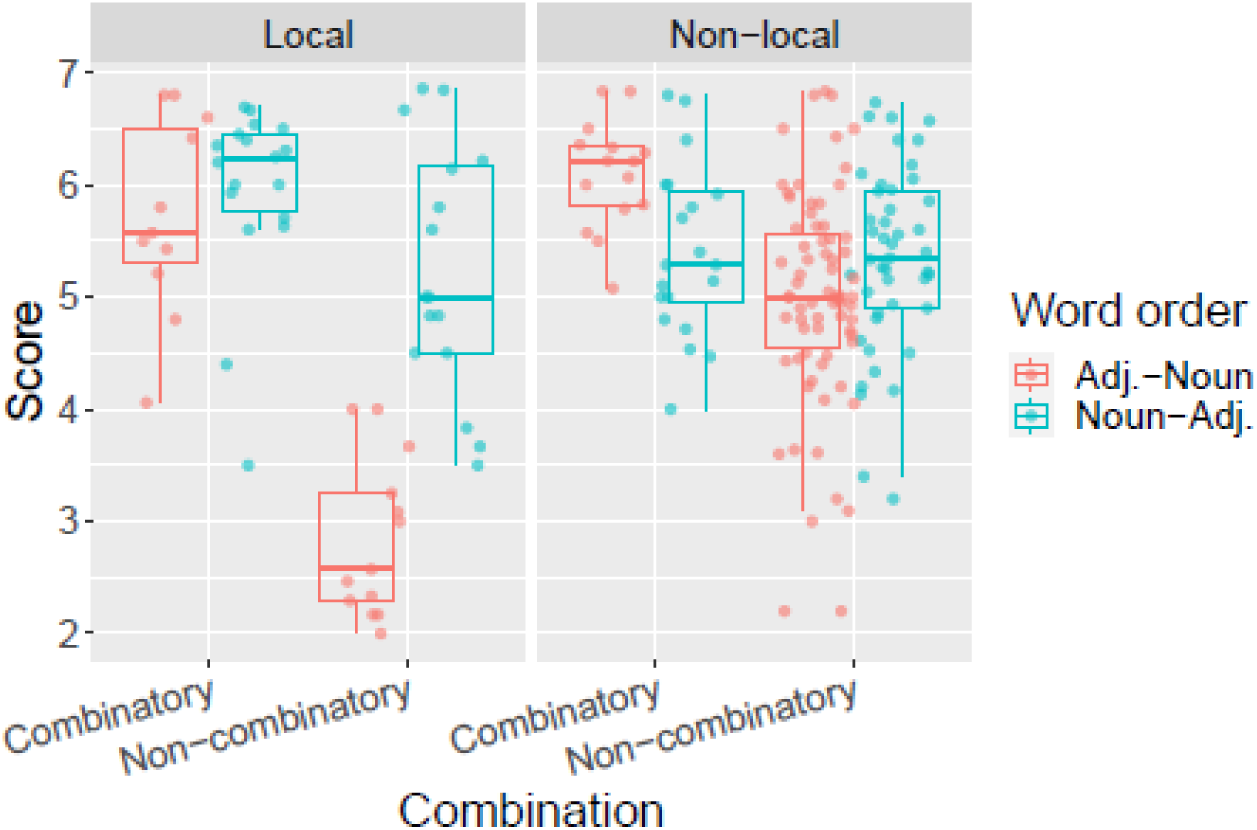
Results from the norming study. The plot shows that, on average, items in the crucial Non-local condition were acceptable. In the Local controls, however, there was a degradation in acceptability for the items in the Non-combinatory Adjective-Noun Word order condition. Each dot represents an individual item that was rated in the study, and the score (y-axis) is the average rating assigned to it from the participants.

**Figure 11:**
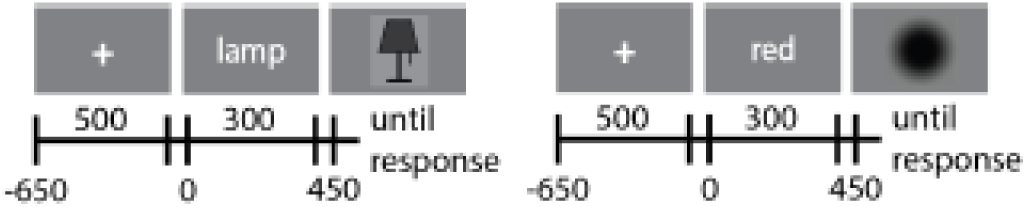
Diagram of the procedure for the single-word picture verification task. The figure shows an example with both nouns and colors. All nouns were presented colored black for this portion of the experiment. In this example the trial on the left should be answered “yes” and the trial on the right should be answered “no.”

**Figure 12:**
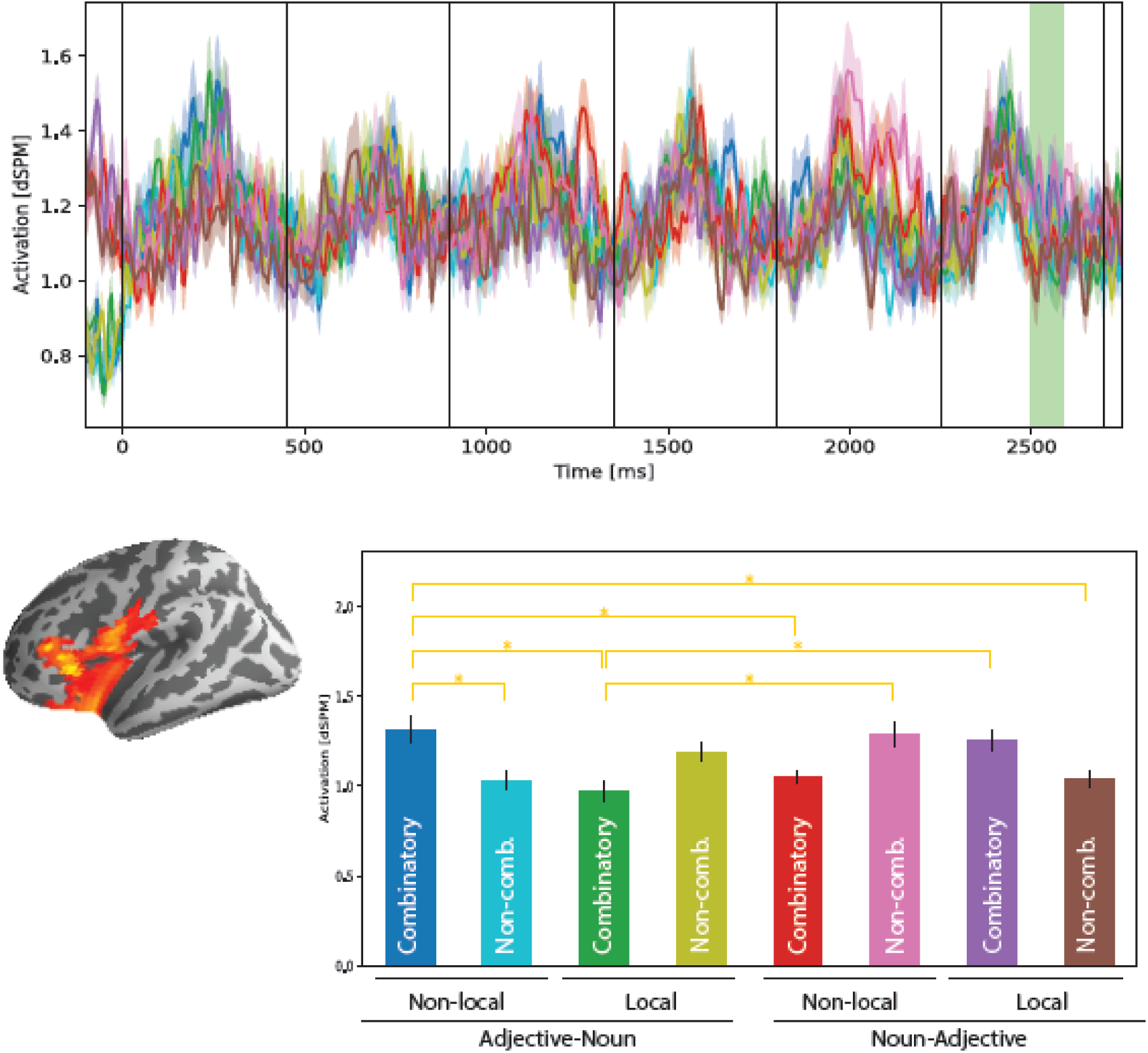
Additional significant results of the spatiotemporal left-hemisphere analysis showing a three-way interaction of Locality by Combination by Word order. The barplot shows the pairwise comparisons that were significant (p < 0.05) after controlling for multiple comparisons.

This step was initially included with the idea that we may be able to use the trials to improve the decoding analysis by combining the single-word trials for a single stimulus word with the relevant condition in the main task. Though we hoped that this step would increase the signal-to-noise ratio of the training data and improve the generalizability of our classifier, it had the opposite effect, and we do not include results with the single-word task in our main results.

## Appendix D. Full 2×2× effect from the spatiotemporal analysis

In addition to the main findings already reported, we also observed a significant 3-way interaction of Locality, Combination, and Word order in the left hemisphere. The significant cluster extended from 250-335ms after target word onset (p=0.03). Spatially, the cluster most strongly localized around the LIFG and into the vmPFC. Note that this cluster is spatially distinct from the one reported in the main text, and likely represents a different aspect of processing.

Figure 12 shows the results of this test. After controlling for multiple comparisons in the pairwise tests, many of the significant results are due to an interaction of factors across the Word order condition, making interpretation of this effect more challenging because differences across Word order conditions in this study were not as well controlled as condition differences within each Word order condition. Overall, we notice that there is an opposite pattern of results within the Adjective-Noun order stimuli and the Noun-Adjective order stimuli, characterized by an increase in activation for the Combinatory stimuli compared to Non-combinatory controls in (i) the Adjective-Noun Non-local condition (p ¡ 0.05) and (ii) the Noun-Adjective Local condition (n.s.), but the opposite pattern in which the Combinatory stimuli had lower activation than the Non-combinatory controls is observed in (i) Adjective-Noun Local condition (n.s.) and (ii) the Noun-Adjective Non-local condition (n.s.). Given that only one of these pairwise comparisons is significant, and given that the interaction is quite complex, it’s very possible that this three-way interaction is not readily interpretable for this experiment.

## Appendix E. Right hemisphere effects observed in the spatiotemporal analysis

In Figure 13, we show the effects observed in the right hemisphere for the same 2×2×2 ANOVA of Combination by Locality by Word order that was described in the main text. We observed a significant interaction of Combination by Word order that, just like in the left hemisphere, was characterized by an increase in activation for the Combinatory condition relative to the Non-combinatory condition for the Adjective-Noun order, but a reversed in the Noun-Adjective order stimuli, where we observed an increase in activation for the Non-combinatory condition. The identified cluster spanned from 150-250ms after target word onset (*p* = 0.004) and mostly localized to the medial wall. As interpretation of effects from the more inner regions of the brain is difficult with MEG, we note that this effect should be interpreted with caution and may not be reliable, especially as previous studies have not reported robust effects along the medial wall for conceptual combination, and it’s possible that due to spatial smearing in the conversion from sensor space to source space, the results shown in Figure 13 are actually picking up effects that originated in the left hemisphere.

**Figure 13:**
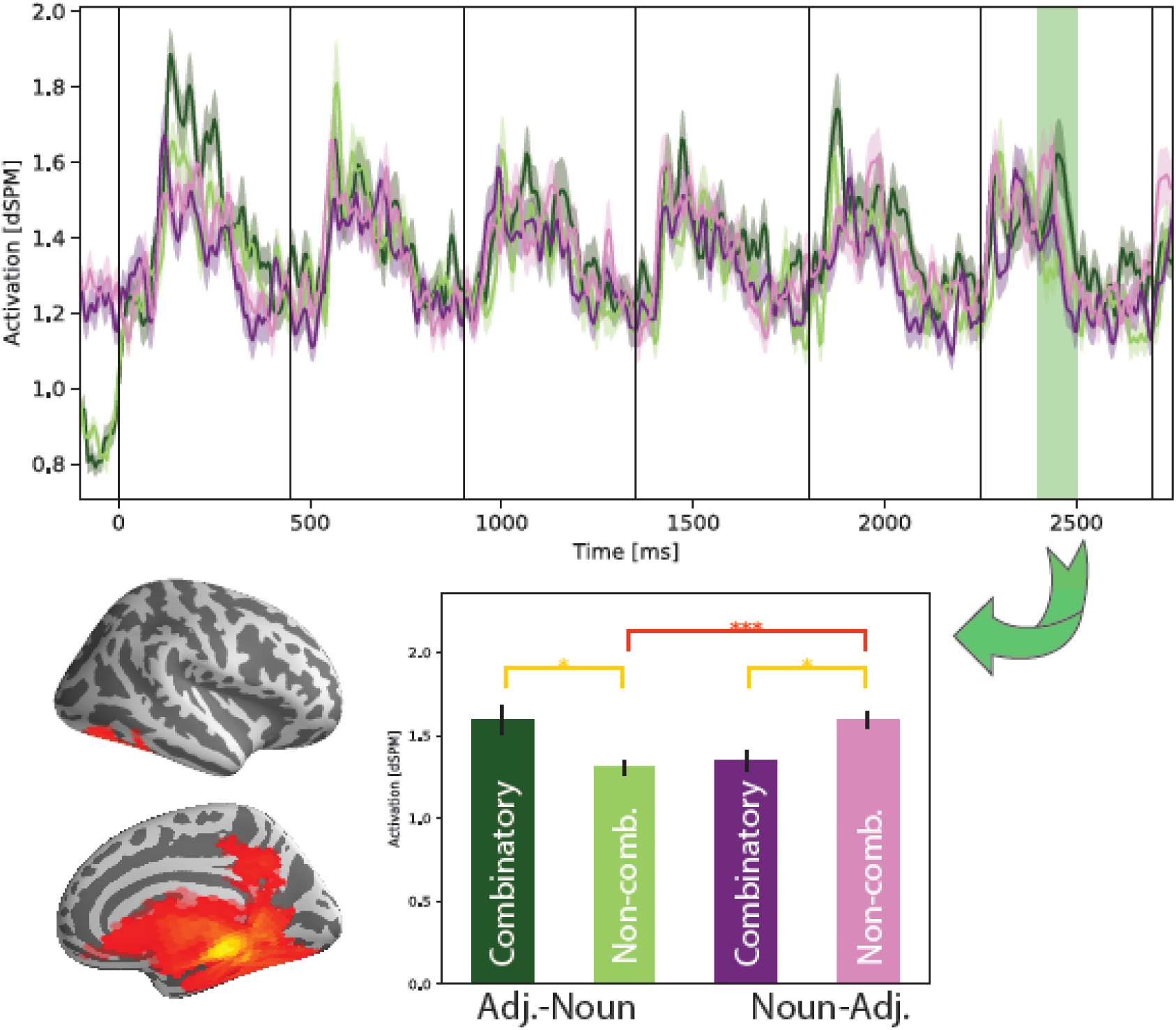
Spatiotemporal 2×2×2 analysis of Word order by Locality by Combination analysis in the right hemisphere showing an analogous pattern of results as was observed in the left hemisphere.

